# Agonist efficiency from concentration-response curves: structural implications and applications

**DOI:** 10.1101/2020.10.13.337410

**Authors:** D. C. Indurthi, A. Auerbach

**Affiliations:** University of Buffalo; SUNY Buffalo

**Keywords:** nicotinic, acetylcholine receptor, dose-response, pharmacology

## Abstract

Agonists are evaluated by a concentration-response curve (CRC), with a midpoint (EC_50_) that indicates potency, a high-concentration asymptote that indicates efficacy and a low-concentration asymptote that indicates constitutive activity. A third agonist attribute, efficiency (η), is the fraction of binding energy that is applied to the conformational change that activates the receptor. We show that η can be calculated from EC_50_ and the asymptotes of a CRC derived from either single-channel or whole-cell responses. For 20 agonists of skeletal muscle nicotinic receptors, the distribution of η values is bimodal with population means at 51% (including ACh, nornicotine and DMPP) and 40% (including epibatidine, varenicline and cytisine). The value of η is related inversely to the size of the agonist’s head-group, with high-versus low-efficiency ligands having an average volume of 70 Å^3^ versus 102 Å^3^. Most binding site mutations have only a small effect on ACh efficiency except for αY190A (35%), αW149A (60%) and those at αG153 (42%). If η is known, the midpoint and high-concentration asymptote can be calculated from each other. Hence, an entire CRC can be estimated from the response to a single agonist concentration, and efficacy can be estimated from EC_50_ of a CRC that has been normalized to 1. Given η, the level of constitutive activity can be estimated from a single CRC.

**Statement of significance:** Receptors are molecular machines that convert chemical energy from agonist binding into mechanical energy of a global conformational change that generates a cell response. Agonists are distinguished by their potency (proportional to affinity) and efficacy, but also by the efficiency at which their binding energy is applied to receptor activation. Here we show that agonist efficiency can be estimated from a single concentration-response curve, and estimate efficiencies of 20 nicotinic receptor agonists. These have a bimodal distribution with larger agonists belonging to the lower-efficiency population. We further show that mutations of some binding site residues alter efficiency, and that knowledge of agonist efficiency simplifies and extends dose-response curve analysis.

## Introduction

Nicotinic acetylcholine receptors (AChRs) are members of the cys-loop, ligand-gated receptor family that in mammals also comprise GABAA, glycine, 5-HT3 and zinc-activated receptors (1). They are 5-subunit, liganded-gated ion channels with agonist binding sites in the extracellular domain, far from a narrow region of the pore in the transmembrane domain that regulates ion conductance (2, 3).

AChRs switch between global C(losed-channel) and O(pen-channel) conformations (‘gating’) to produce transient membrane currents. Agonists promote channel opening because they bind more strongly to the O conformation. Importantly, the energy (structure) of the binding site at the gating transition state resembles that of O (4). Hence, when a receptor begins its journey from C to O, extra (favorable) binding energy eases the pathway, thereby increasing the probability of reaching and residing in O (P_O_) (5, 6).

AChRs are the primary receptors at vertebrate neuromuscular synapses where they initiate muscle membrane depolarization and contraction. Neuromuscular AChRs have two α1 subunits and one each of β, δ and either ε (adult) or γ (fetal). There are two neurotransmitter binding-sites located at α1–δ and α1–ε/γ subunit interfaces (7) that are approximately equivalent for ACh in adult-type AChRs (5). AChRs switch conformation spontaneously (only under the influence of temperature), with the presence of neurotransmitters at both adult sites increasing the opening rate constant by a factor of ~5 million and the lifetime of the O conformation by a factor of ~5.

Agonists are typically characterized by a potency (proportional to affinity) and an efficacy. Affinity is a measure of how strongly the ligand binds to its target site and is the inverse of an equilibrium dissociation constant. The constants K_dC_ and K_dO_ correspond to low-affinity (weak) binding to the C conformation and high-affinity (strong) binding to the O conformation. (The logarithm of an equilibrium dissociation constant is proportional to binding energy.)

The high-concentration asymptote of a CRC, or the maximum response elicited by the ligand, is called P_O^max^_ in single-channel or I^max^ in whole-cell currents. This limit gives the agonist’s efficacy and depends only on the fully-liganded gating equilibrium constant. The midpoint of a CRC, or the agonist concentration that produces a half-maximal response is proportional to K_dC_ but also depends on the gating equilibrium constant.

The low-concentration asymptote of a CRC, which gives the level of activity in the absence of agonists (P_O_^min^ or I^min^), depends on the unliganded gating equilibrium constant that is typically small and difficult to measure. However, it is important to know the exact value of this constant because it multiplies the fully-liganded gating equilibrium constant to influence potency, efficacy and synaptic current profiles. Allosteric modulators and AChR mutations (8), including some that cause slow-channel myasthenic syndromes (9), alter EC_50_, I^max^ and the time course of synaptic currents simply by increasing or decreasing the unliganded gating equilibrium constant, without making a noticeable change in baseline activity.

Recently, efficiency (η; eta) was defined as the fraction of an agonist’s chemical binding energy that is converted into the mechanical (kinetic) energy for gating (10). Efficiency reports the strength of the link between binding and gating. As shown previously (and again below by using a different approach), η is a function of the resting/active binding energy ratio, logK_dC_/logK_dO_. Direct, independent measurements of these two equilibrium dissociation constants obtained by detailed kinetic modeling of single-channel currents indicated that at adult-type human AChR neurotransmitter binding sites, ACh and 3 related agonists on average apply ~50% of their binding energy to gating whereas at the α1–δ binding site the frog toxin epibatidine and 3 related agonists on average apply only ~40% (10).

Here, we show that agonist efficiency can be estimated from the asymptotes and midpoint of a single CRC constructed from either single-channel or whole-cell responses. Given two agonists with the same EC_50_, the one with the larger I^max^ has the greater η. We provide separate efficiency estimates for 20 agonists of mouse adult AChRs and show that knowledge of agonist efficiency broadens our understanding of receptor activation and drug action.

## Materials and Methods

### Experimental design

#### Expression

Human embryonic kidney (HEK) 293 cells were maintained in Dulbecco’s minimal essential medium supplemented with 10% fetal bovine serum and 1% penicillin–streptomycin, pH 7.4. For single-channel experiments, AChRs were expressed by transient transfection of 3 μg mouse α1, β, δ and ε subunits in the ratio 2:1:1:1 (*TransIT*^®^ 293 transfection reagent; Mirus Bio, Madison, WI). Electrophysiological experiments started ~48 hours post-transfection. For whole cell recording, HEK-293 cells were transiently transfected with adult-type mouse AChRs using calcium phosphate precipitation. 20 μg of cDNA was added in the ratio of 2:1:1:1 (α1-GFP encoded between M3-M4, β, δ and ε) to a T75 flask at ~60% confluence. Cells were incubated for ~16 hr at 37° C, replenished with fresh medium and harvested after ~20 hrs of washing. GFP positive cells were sorted by using a ABD FACS Fusion 4 laser Cell Sorter (Becton Dickinson, Franklin Lakes, NJ). Cells were excited with laser at 488 nm and the GFP signal was collected in the green channel through a 530/40 filter. A light scatter gate was drawn in the SSC versus FS plot to exclude debris and to include viable single cells. No animals were used in this study.

#### Electrophysiology

Single-channel currents were recorded in the cell-attached patch configuration at 23° C. The bath solution was (in mM): 142 KCl, 5.4 NaCl, 1.8 CaCl_2_, 1.7 MgCl_2_, 10 HEPES/KOH (pH 7.4). Patch pipettes were fabricated from borosilicate glass and fire polished to a resistance of ~10 MΩ when filled with the pipette solution that was Dulbecco’s phosphate-buffered saline (in mM): 137 NaCl, 0.9 CaCl_2_, 2.7 KCl, 1.5 KH_2_PO_4_, 0.5 MgCl_2_, and 8.1 Na_2_HPO_4_ (pH 7.3/NaOH). Currents were recorded using a PC505 amplifier (Warner instruments, Hamden, CT), low-pass filtered at 20 kHz and digitized at a sampling frequency of 50 kHz using a data acquisition board (SCB-68, National instruments, Austin, TX). Agonists were added to the pipette solution at the desired concentration.

Whole-cell currents were recorded using an IonFlux 16 automated patch-clamp system (Fluxion, California, USA) on 96-well IonFlux microfluidic ensemble plates that give a cumulative whole-cell current from up to 20 cells. GFP-positive cells were re-suspended in extracellular solution containing (in mM): 138 NaCl, 4 KCl, 1.8 CaCl_2_, 1 MgCl_2_, 5.6 glucose and 10 HEPES, pH adjusted to 7.4 with NaOH. Cells were captured in the trapping wells with intracellular solution containing 60 KCl, 70 KF, 15 NaCl, 5 HEPES and 5 EGTA, pH adjusted to 7.2 using KOH. Cells clamped at −80 mV were exposed to a 2 s agonist application followed by 90 s wash between applications to allow recovery from desensitization. IonFlux software (ver.4.5) was used for cell capture, seal formation, compound application and data acquisition.

#### Analysis

Scheme 1 (Fig. 1) shows the main states of AChR activation/de-activation. First, we estimated η from single-channel current CRCs. When agonist-binding and channel-opening rate constants are sufficiently large, openings occur in clusters (11) (Fig. 2A, Fig. S1-S4). Shut intervals within clusters represent mainly agonist binding to C and channel opening (bold in Fig. 1), whereas shut intervals between clusters represent mainly long-lived desensitization (not shown in Fig. 1; for connections see (6)). We selected for analysis clusters that appeared by eye to arise from a homogeneous P_O_ population and, in order to exclude sojourns in desensitized states, limited our analyses to intra-cluster interval durations.

**Figure 1.**
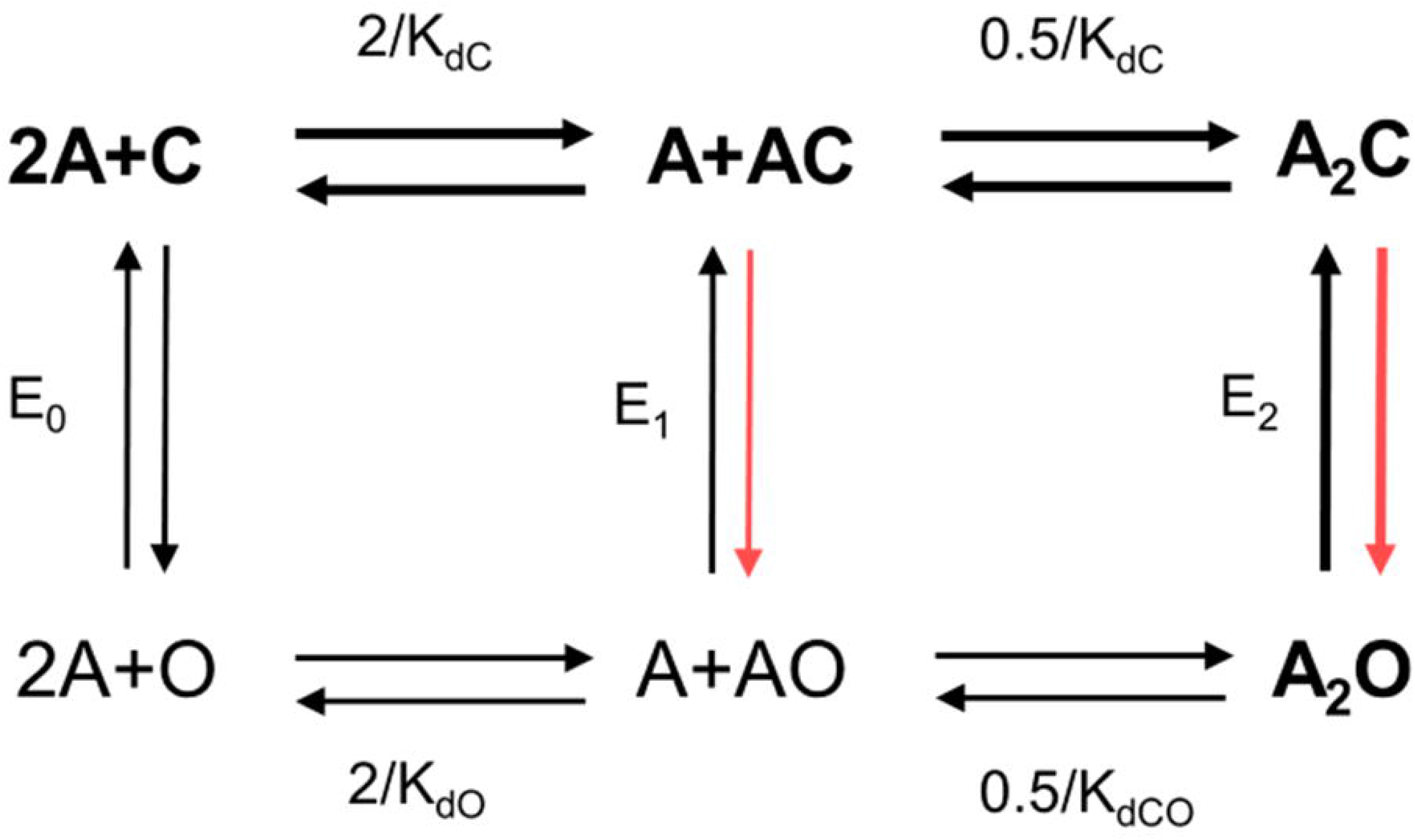
Cyclic activation of AChRs (Scheme 1). Receptors switch between C(losed-channel) and O(pen-channel) conformations spontaneously (influenced only by temperature) with or without agonists (A). Equilibrium constants: E_n_, gating with n bound agonists; K_dC_ and K_dO_, dissociation constants to C (low affinity) and to O (high affinity). The adult-type, endplate AChR binding sites are approximately equivalent and independent with regards to the agonists used in this study. From experiments and microscopic reversibility, E_2_/E_0_=(K_dC_/K_dO_)^2^. Agonists increase activity above the baseline level because they bind more-strongly to the C-O transition state, with the extra binding energy serving to increase the channel-opening rate constant (red arrows). Thick arrows and bold letters mark the physiological activation-deactivation pathway.

**Figure 2.**
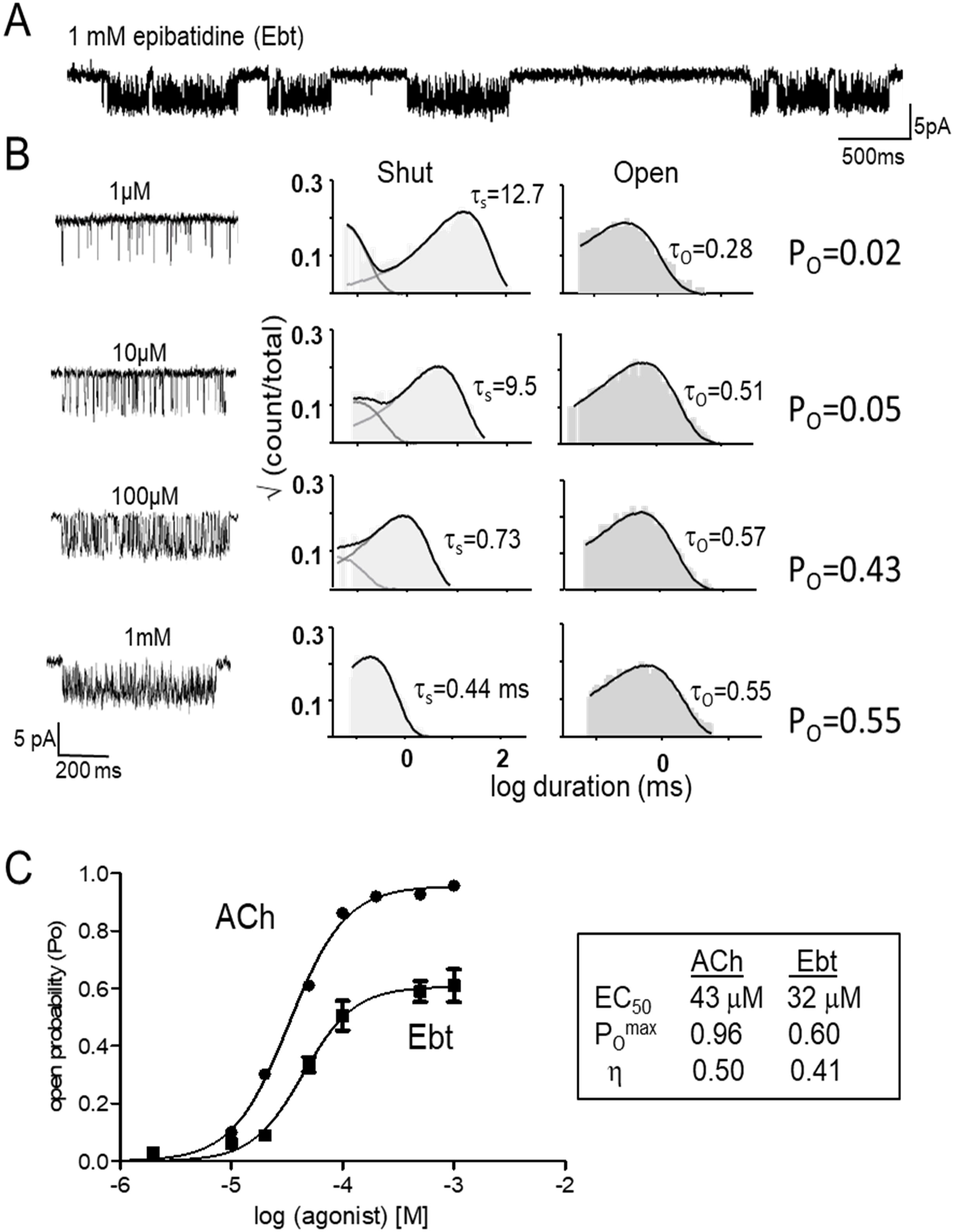
Single-channel current CRCs. A. Low-resolution view of cell-attached, single-channel currents activated by epibatidine (Ebt; open is down). Clusters of open/shut intervals from one AChR separated by silent periods in which all AChRs in the patch are desensitized. B. Left, high-resolution views of example clusters at different Ebt concentrations. Right, corresponding intra-cluster interval-duration histograms. τ_s_ and τ_o_, predominant shut- and open-interval time constants (in ms). Open-channel probability (P_O_) at each agonist concentration is τ_o_/(τ_s_+τ_o_). C. Each CRC was fitted by Eq. 1 to estimate EC_50_ and P_O^max^_ (symbols, mean±s.e.m.). Efficiency (η) was calculated by using Eqs. 2–5. It is apparent that ACh is more efficient than Ebt because the same EC_50_ is associated with a greater P_O^max^_.

Because of the high extracellular [K^+^], the cell membrane potential (V_m_) was 0 mV. The AChR agonists we examined also are channel-blockers. To both generate measurable currents and reduce the effect of channel block on PO, the membrane was depolarized to +70 mV by holding the pipette at −70 mV. This effectively eliminated agonist binding to the channel-block site in the transmembrane domain but did not affect agonist binding to the neurotransmitter sites in the extracellular domain.

Analyses of the single-channel (outward) currents were performed by using QUB software (12). A cluster was defined as a group of openings flanked by shut intervals longer than a duration that depended on the agonist concentration (range, 7-20 ms). Open and shut currents within clusters were idealized into noise-free intervals by using the segmental k-means algorithm after digitally low-pass filtering the data at 10 kHz (13). Idealized interval durations were fitted by multiple exponential components using a maximum interval likelihood algorithm (14). Cluster P_O_ at each agonist concentration was calculated from the time constants of the predominant components of the shut- (τ_s_) and open-time distributions (τ_o_): τ_o_/(τ_s_+τ_o_) (Fig. 2B). The single-channel CRC was a plot of the absolute P_O_ (not normalized) versus the agonist concentration.

We also estimated η from whole-cell current CRCs. The currents were digitized using a sampling frequency of 10□kHz and were analyzed using IonFlux Data Analyzer v5.0. Peak currents were normalized to a maximum response (I/I^max^), where I^max^ was the response to 300 μM ACh. The 20-80% rise time to a step to 300 μM ACh was ~400 ms, a time we attribute to solution exchange.

The rate of entering a long-lived desensitized state is proportional to cluster P_O_ and occurs with a rate constant of ~5 s^−1^ (15). Hence, under conditions where P_O_ is ~1, a whole-cell current will decline with a time constant of ~200 ms. As a consequence, the peaks of whole-cell currents elicited by high-concentrations of high-efficacy agonists are truncated because of the solution exchange time. This has the effect of shifting EC_50_ to lower concentrations. Responses at lower agonist concentrations or from lower-efficacy agonists were unaffected by desensitization.

#### Voltage, E_0_ and background mutations

Depolarization to V_m_=+70 mV reduces channel block by the agonist but has the undesired consequence of shortening τ_o_ to make single-channel current detection and idealization difficult. To compensate, we added the background mutation εS450W (in the M4 transmembrane segment of the ε subunit) that has the equal-but-opposite effect on the unliganded gating equilibrium constant E_0_ as does depolarization by +140 mV, but has no effect on agonist binding (16). With this mutation, τ_o_ and E_0_ at +70 mV were the same as in wild-type (wt) adult AChRs at V_m_=−70 mV. E_0_ at −100 mV is 7.4 x 10^−7^ and is reduced e-fold by a 60 mV depolarization (17). Hence, we estimate that in our experiments at V_m_=+70 mV and with εS450W, E_0_ was 5.2 x 10^−7^. In the whole-cell experiments, no background mutations were used and V_m_=−80 mV, so we estimate E_0_ was 5.9×10^−7^.

With the low-efficacy agonists varenicline, tetraethylammonium and tetramethylphosphonium, single-channel clusters generated by wt AChRs were poorly defined because the channel-opening rate constant was small. For these ligands, P_O_ could not be estimated accurately using wt AChRs. To increase the diliganded opening rate constant and generate better-defined, higher-P_O_ clusters, we added two background mutations in the ε subunit, εL269F (in the M2 helix) and εE181W (in strand β9), without εS450W. Together, these two substitutions increase E_0_ by 1084-fold (making it 4.9×10^−4^) without affecting agonist binding (18, 19). From the uncorrected CRC, we estimated an E_2_ value from the P_O^max^_ (Eq. 4) and K_dC_ from EC_50_. We divided this E_2_ by 1084 to arrive at a corrected E_2_ from which we calculated corrected P_O^max^_ and EC_50_ values that pertain to wt AChRs (Table 1).

**Table 1.** Efficiencies from single-channel CRCs. EC_50_, P_O^max^_ and n_H_ obtained by using Eq. 1. n, number of CRCs; sem, standard error of the mean. E_2_, diliganded gating equilibrium constant; K_dC_, equilibrium dissociation constant to C; K_dO_, equilibrium dissociation constant to O (see Fig. 1). The unliganded gating equilibrium constant E_0_ was 5.2×10^−7^. η, efficiency calculated using Eq. 5; volume is of the agonist’s head-group volume (Fig. S5). Previously published values are from ^□^(23), ^±^(28), ^ϕ^(5), ^Ψ^ (29), ^□^(10). All entries pertain to wt adult AChRs.

| Measured Values |  |  |  |  |  | Calculated values |  |  |  |  |  |  |
| --- | --- | --- | --- | --- | --- | --- | --- | --- | --- | --- | --- | --- |
| s.no | Agonist | EC<sub>50</sub>±sem<br>(μM) | P<sub>O</sub><sup>max</sup>±sem | n<sub>H</sub> | n | E<sub>2</sub> | K<sub>dC</sub><br>(μM) | K<sub>dO</sub><br>(nM) | log K<sub>dC</sub> | log K<sub>dO</sub> | □ | volume<br>(Å<sup>3</sup>) |
| 1 | ACh<sup>□</sup> | 43 | 0.96 | 1.7 | - | 23.4 | 174 | 24 | -3.76 | -7.62 | **0.50** | 77 |
|  | ACh<sup>Φ</sup> |  |  |  |  |  | 130 | 18 | -3.89 | -7.74 | **0.50** |  |
|  | ACh<sup>Ψ</sup> | 22 | 0.96 |  |  | 24 | 90 | 12 | -4.04 | -7.91 | **0.49** |  |
| 2 | Nor<sup>±</sup> | 72 | 0.92 | 1.0 | - | 12 | 200 | 38 | -3.70 | -7.41 | **0.50** | 69 |
| 3 | CCh<sup>□</sup> | 320 | 0.84 | 1.6 | - | 5.25 | 542 | 159 | -3.27 | -6.80 | **0.52** | 77 |
| 4 | Ana<sup>±</sup> | 524 | 0.78 | 1.2 | - | 3.63 | 719 | 253 | -3.14 | -6.60 | **0.52** | 70 |
| 5 | TMA<sup>□</sup> | 1200 | 0.75 | 1.1 | - | 3 | 1480 | 574 | -2.83 | -6.24 | **0.54** | 77 |
| 6 | DMT<sup>□</sup> | 4200 | 0.42 | 0.8 | - | 0.72 | 2730 | 2150 | -2.56 | -5.67 | **0.54** | 58 |
| 7 | DMP<sup>□</sup> | 6700 | 0.26 | 1.1 | - | 0.35 | 3570 | 4040 | -2.45 | -5.39 | **0.54** | 56 |
| 8 | Cho | 4013 | 0.05 |  | - | 0.05 | 4100 | 15100 | -2.39 | -4.82 | **0.50** | 77 |
| 9 | DMPP<sup>\$</sup> | 246±81 | 0.87±0.09 | 1.3 | 3 | 6.69 | 480 | 134 | -3.32 | -6.87 | **0.52** | 59 |
| 10 | 4OH-B<sup>\$</sup> | 2171±77 | 0.29±0.04 | 0.9 | 3 | 0.41 | 1270 | 1350 | -2.92 | -5.87 | **0.50** | 77 |
| 11 | 3OH-P<sup>\$</sup> | 3485±10 | 0.15±0.02 | 0.6 | 4 | 0.18 | 1660 | 2840 | -2.78 | -5.55 | **0.50** | 77 |
| 12 | Ebt<sup>\$</sup> | 32±2 | 0.60±0.6 | 1.9 | 4 | 1.50 | 28 | 16 | -4.55 | -7.78 | **0.41** | 88 |
| 13 | Ebx<sup>\$</sup> | 90±4 | 0.74±0.02 | 1.6 | 3 | 2.85 | 108 | 46 | -3.96 | -7.33 | **0.46** | 88 |
| 14 | Cyt<sup>\$</sup> | 137±5 | 0.18±0.01 | 1.3 | 4 | 0.22 | 67 | 104 | -4.17 | -6.99 | **0.40** | 114 |
| 15 | Var<sup>*</sup> | 135±12 | 0.015±0.002 | 4.8 | 3 | 0.02 | 57 | 331 | -4.25 | -6.48 | **0.35** | 102 |
| 16 | TEA<sup>*</sup> | 4200±130 | 0.002±0.0007 | 1.1 | 3 | 0.002 | 174 | 26800 | -2.76 | -4.57 | **0.40** | 136 |
| 17 | TMP<sup>*</sup> | 844±32 | 0.03±0.002 | 2.5 | 3 | 0.031 | 360 | 1473 | -3.44 | -5.83 | **0.41** | 87 |
| 18 | Atx<sup>□</sup> |  |  |  |  |  | 115 | 247 | -3.94 | -6.61 | **0.40** | 114 |
| 19 | Aza<sup>□</sup> |  |  |  |  |  | 934 | 6053 | -3.03 | -5.22 | **0.42** | 88 |
| 20 | Nic<sup>±</sup> |  |  |  |  | 0.87 | 1000 | 920 | -3.00 | -6.04 | **0.50** | 84 |

#### Equations

Single-channel CRCs were constructed from P_O_ values after eliminating extraneous events arising from channel block, desensitization and modal activity (20). Whole-cell CRCs were constructed directly from peak currents. EC_50_ and P_O^max^_ (or I^max^) the Hill coefficient (n_H_) were estimating by fitting the CRC,

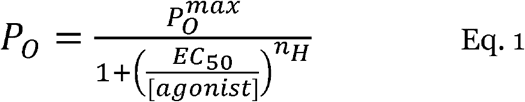

Scheme 1 (Fig. 1) was used to derive expressions for η. Because microscopic reversibility is satisfied,

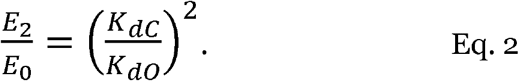

E_2_ and E_0_ are the diliganded and unliganded gating equilibrium constants and K_dC_ and K_dO_ are the equilibrium dissociation constants for binding to C and O. The exponent reflects the fact that in adult-type AChRs there are 2 neurotransmitter sites that are approximately equivalent and independent with regards to the agonists used in this study. Eq. 2 has been confirmed by experiment (5).

Constitutive and mono-liganded activity are both rare, so in wt AChRs the only significant pathway that generates current is the clockwise, linear activation route highlighted in Fig. 1. Transitions between these 4 states determine P_O_ and, hence, the experimental values of EC_50_ and P_O^max^_ (or I^max^).

Accordingly, EC_50_ depends on both binding and gating equilibrium constants,

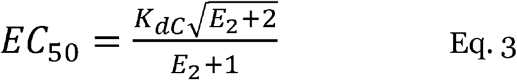

whereas P_O^max^_ (I^max^) depends only on the gating equilibrium constant,

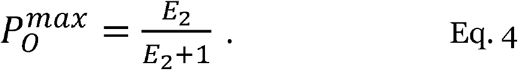

This equation also can be used to relate P_O^min^_ (I^min^) and E_0_.

Agonist efficacy depends on the diliganded gating equilibrium constant that from Eq. 2 is a function of the affinity ratio, K_dC_/K_dO_. Taking the log of Eq. 2, we see that efficacy is determined by the *difference* between the binding energies, log(K_dC_)-log(K_dO_). Partial agonists experience smaller increases in O versus C binding energy compared to full agonists, antagonists experience no change in binding energy, and inverse agonists experience a decrease in favorable binding energy upon receptor activation.

In contrast, η depends on the *ratio* of these binding energies, log(K_dC_)/log(K_dO_), as shown previously (10) and as follows. Agonist activation of a resting, unliganded AChR entails connecting the resting-unliganded state C to the diliganded-active state A2O (Fig. 1). The product of the equilibrium constants (or sum of the energy changes) for steps linking these states in the clockwise direction (the highlighted, physiological activation route) is the same as in the rarely-taken, counter-clockwise direction. The product of the counter-clockwise constants is E_0_/K_dO^2^_, the negative log of which is proportional to the *total* energy required for constitutive gating and binding to O at 2 sites, 2log(K_dO_)-logE_0_. The product of the equilibrium dissociation constants connecting C with A2C is 1/K_dC^2^_, the negative log of which is proportional to the energy for just the *binding* part of clockwise activation, 2log(K_dC_). We are interested only in the agonist component of the total energy and, because E_0_ is agonist-independent, it can be ignored. Hence, the fraction of the *total* agonist energy that is used in *binding* is 2logK_dC_/2logK_dO_, so efficiency, or the fraction of this total that is applied to *gating*, is

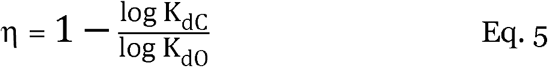

Efficacy and efficiency are distinct, but related, agonist attributes. In terms of energy, efficiency is equal to efficacy (logK_dO_-logK_dC_) divided by logK_dO_ (Eq. 5). An agonist can be high efficacy and low efficiency (epibatidine) or low efficacy and high efficiency (choline), but within limits. If an agonist has P_O^max^_=0.75 (about the same as TMA) and η=30% it would have unreasonably small equilibrium dissociation constants, K_dC_=27 nM and K_dO_=15 pM. In practice, high-efficacy agonists will also have high efficiencies.

Except for Fig 8A, we calculated efficiency from EC_50_, P_O^max^_ and P_O^min^_ using a stepwise approach: i) E_2_ from P_O^max^_ (Eq. 4), ii) K_dC_ from E_2_ and EC_50_ (Eq. 3), iii) K_dO_ from E_2_ and K_dC_ using a known value of E_0_ (Eq. 2), and, finally, iv) η from the equilibrium dissociation constant ratio (Eq. 5). In Fig. 8A only, an approximate value of η was calculated directly using Eq. 10 with A=0.

Four prior results enabled us to estimate η from a CRC. First, adult AChR binding sites have approximately the same affinity, so only single values of the equilibrium dissociation constants needed to be estimated for each ligand. Second, Scheme 1 and microscopic reversibility have been proved experimentally. Third, the unliganded gating equilibrium constant has been measured. In an ‘efficiency’ plot for a group of ligands (10), E_0_ is estimated from the y-intercept (see Eq. 8, below). However, prior knowledge of I^min^ (~E_0_) is required to estimate efficiency from a single CRC. I^min^ is the same for all agonists and so needs to be estimated only once for each receptor (at a given membrane potential).

#### Statistical analyses

For both single-channel and whole-cell CRCs, the midpoint, maximum and slope (EC_50_, P_O^max^_ or I^max^, and n_H_) were estimated by fitting by Eq. 1 using GraphPad Prism 6 (GraphPad Inc., San Diego, CA). Eq. 9 was solved numerically for E_2_ using the symbolic math program Wolfram Alpha.

The goodness of fit for the efficiency frequency distribution (Fig. 5A) was estimated using Prism. The F-test rejects the null hypothesis (Gaussian fit) over sum of two Gaussian with an F-value (F=3.9) and significance (P-value, <0.05). A k-means cluster analysis algorithm (MATLAB^®^, MathWorks, Natick, MA) was used to define agonist groups for 2D cluster analysis, both efficiency and head-group volume (Fig. 5B). Correlation significance between log EC_50_ or log P_O^max^_, measured from CRCs or calculated from each other (Fig. 8A) was by Pearson’s correlation test, using Prism software. The P-value (two-tail) <0.0001 and r^2^=0.78 or 0.74 imply that there is a significant correlation.

#### Agonists

Agonist structures are shown in Fig. 3, Fig. 4 and Fig. S5. Agonist head-group volumes (Fig. 5B) were calculated using Chimera (21). Acetylcholine (ACh), nornicotine (Nor), nicoztinc (Nic), carbamylcholine (CCh), anabasine (Ana), tetramethylammonium (TMA), dimethylthiazolidinium (DMT), dimethylpyrrolidium (DMP), choline (Cho), 3-hydroxypropyltrimethylammonium (3OH-PTMA), 4-hydroxybutyltrimethylammonium (4OH-BTMA), anatoxin (Atx), azabicycloheptane (Aza), tetraethylammonium (TEA), epibatidine (Ebt), epiboxidine (Ebx), varenicline (Var), cytisine (Cyt), dimethylphenylpiperazinium (DMPP), tetramethylphosphonium (TMP). Cyt, Var, TEA and TMP were from Sigma^®^ (St. Louis, MO). The sources for other agonists are given in previous publications (10, 22).

**Figure 3.**
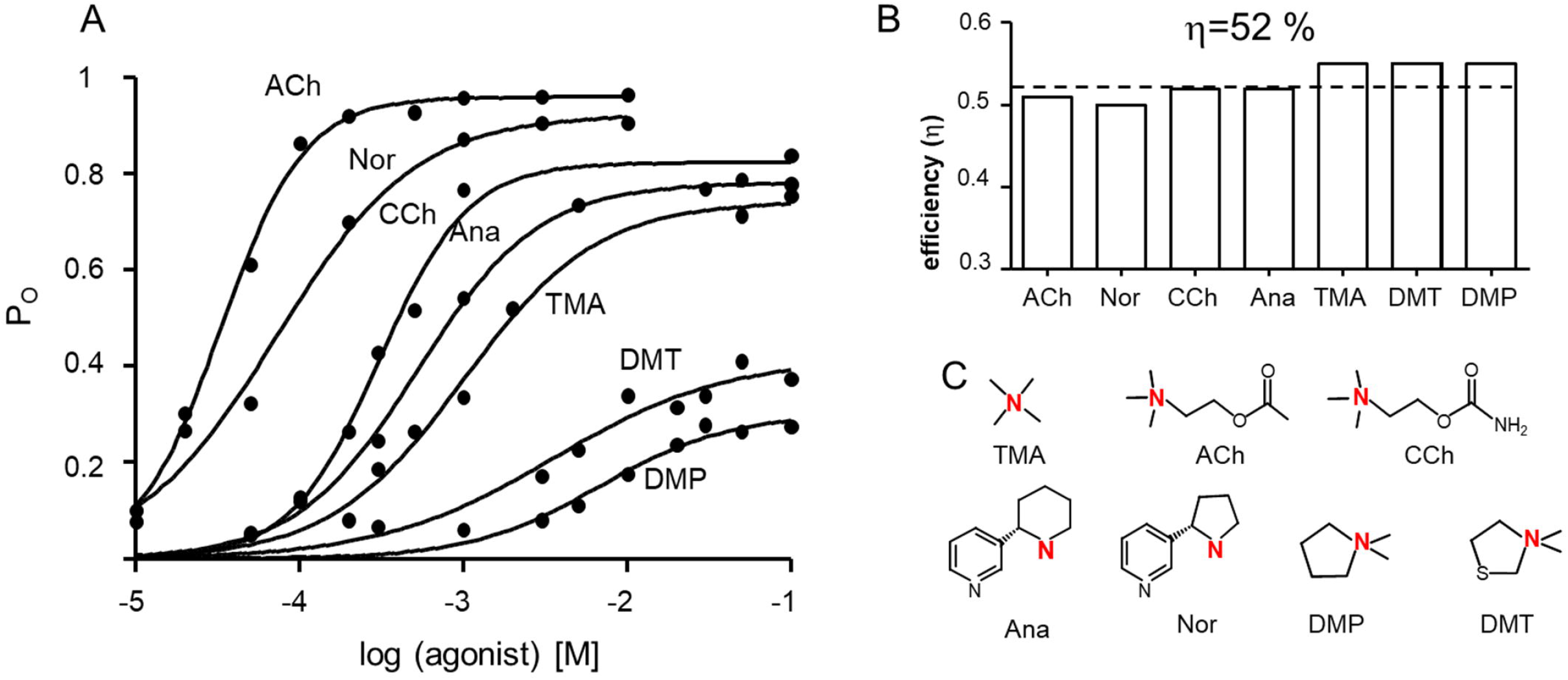
Efficiencies from single-channel CRCs. A. CRCs of 7 agonists in adulttype mouse AChRs (re-plotted from (23)). There is an inverse correlation between EC_50_ and P_O^max^_ (Table 1). B. Agonist efficiencies calculated from EC_50_ and P_O^max^_. All agonists have a similar efficiency (average, 52%; dashed line). C. Agonist structures (see Materials and Methods for abbreviations). Red, key nitrogen atom in the agonist’s head-group.

**Figure 4.**
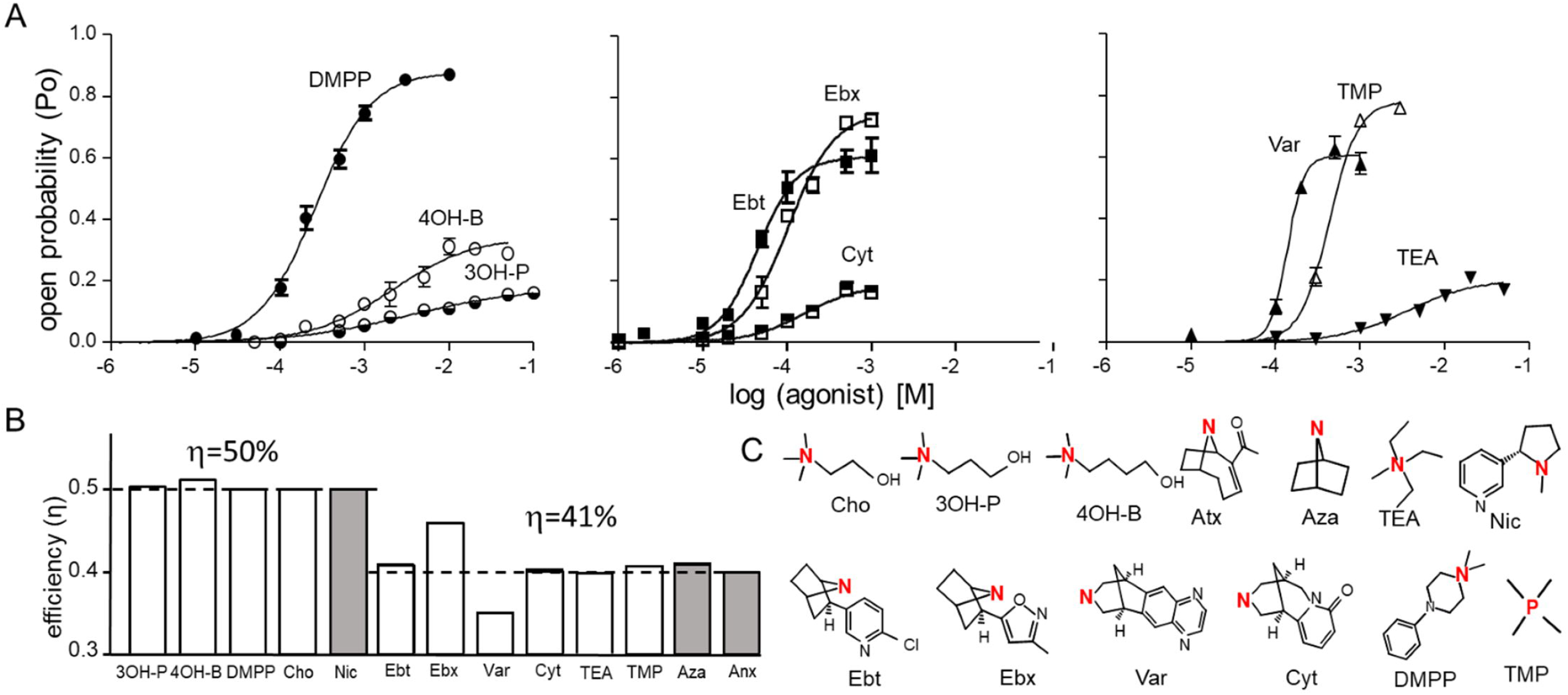
Efficiencies from more single-channel CRCs. A. Single-channel CRCs of 9 agonists in adult-type mouse AChRs (symbols, mean±s.e.m.). In some cases background mutations were used to increase constitutive gating and, hence, increase P_O^max^_ and left-shift EC_50_ (εS450W, left and middle; εL269F+εE181W, right). EC_50_ and P_O^max^_ values in Table 1 have been corrected for these backgrounds and pertain to wt AChRs. B. Efficiencies calculated from the CRCs (open bars) or from previously reported measurements of K_dC_ and K_dO_ obtained by kinetic modeling (gray bars; (5, 28)). There are 2 populations with average efficiencies of 50% and 41% (dashed lines). C. Agonist structures. Red, key nitrogen or phosphorous atom in the agonist’s head-group.

**Figure 5.**
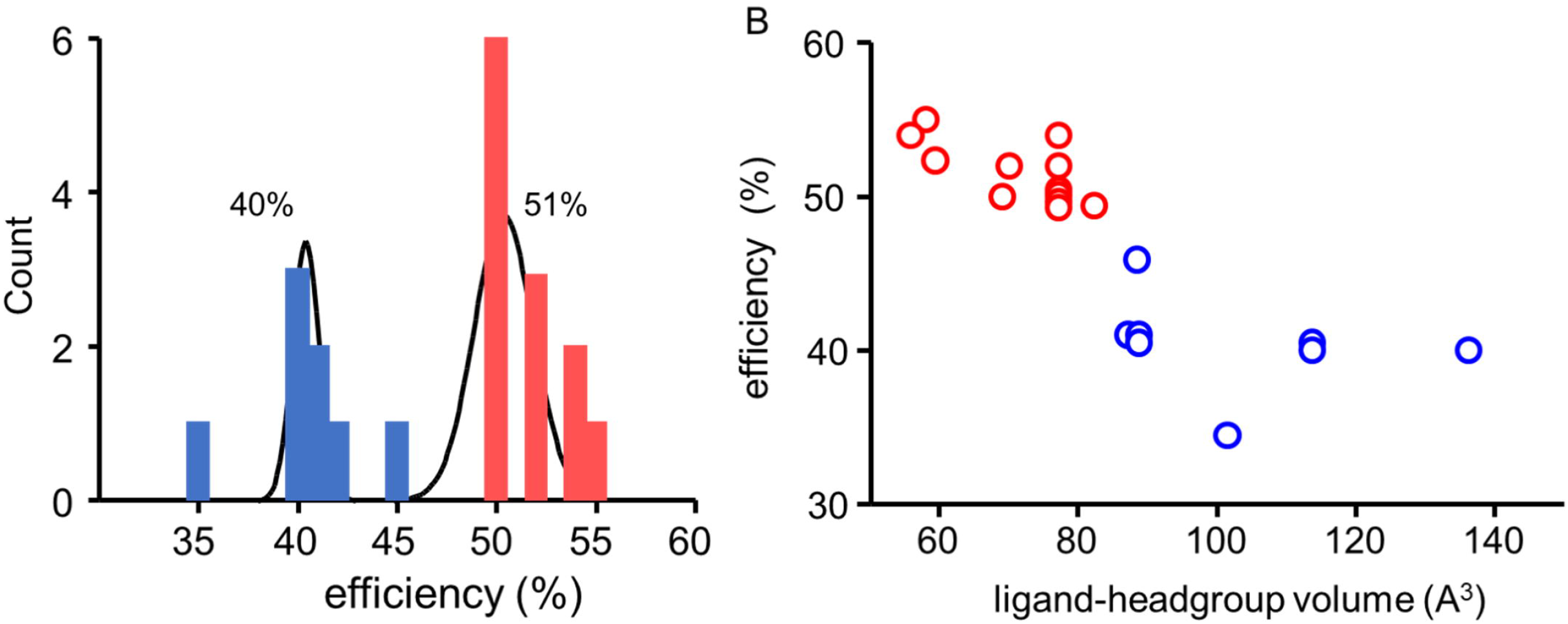
Distribution of agonist efficiency. A. Distribution of efficiencies for 20 agonists, fitted by the sum of two Gaussians. There are 2 populations at η=51±2 % and 40±4 % (mean±s.d.). B. 2-D scatter plot of efficiency versus head-group volume (v). Cluster analysis (k-means) shows that there are 2 populations with η/v centroids at 52%/70.4 Å^3^ (red) and 41%/102.2 Å^3^ (blue).

## Results

### ACh efficiency

Fig. 2 shows example single-channel currents and CRCs. For the neurotransmitter ACh, P_O^max^_ and EC_50_ estimated by fitting the CRC by Eq. 1 are 0.96 and 43 μM (Table 1). From P_O^max^_ we calculate the diliganded gating equilibrium constant, E_2_=23.4 (Eq. 4). From this value and EC_50_ we calculate the low-affinity equilibrium dissociation constant, K_dC_=174 μM (Eq. 3). The unliganded gating equilibrium constant at +70 mV is 5.2×10^−7^ (see Methods) so we calculate the high-affinity equilibrium dissociation constant is K_dO_=26 nM (Eq. 2). Finally, from the ratio of the logs of the two equilibrium dissociation constants we calculate the efficiency of the neurotransmitter, η_ACh_=50% (Eq. 5).

We also calculated η_ACh_ from published values of K_dC_ and K_dO_ obtained either from wild-type mouse AChRs (23) or from individual α-δ and α-ε human AChR binding sites (5), in both instances estimated by kinetic modeling of single-channel currents. The efficiencies calculated from these independent datasets are both 50% (Table 1).

At adult AChR binding sites, half of the neurotransmitter binding energy is applied to the gating conformational change. That is, at each of the 2 binding sites the energy change when ACh binds to the C conformation is approximately equal to the increase in binding energy that happens within the C-to-O transition.

### Efficiency of other agonists

We fitted other previously-published, single-channel CRCs (23) to estimate EC_50_ and P_O^max^_, and from these calculated agonist η values as described above (Fig. 3 and Table 1). Despite the wide ranges in both EC_50_ (43 μM to 6.7 mM) and P_O^max^_ (0.26 to 0.96), all 8 of these agonists (including ACh) have a similar efficiency, η=52+2% (mean+s.d.) (Fig. 3B). The efficiency of the lowest-efficacy agonist in this group, DMP, was greater than that of the highest-efficacy agonist, ACh. This highlights the distinction between efficiency (that depends on the binding energy ratio) and efficacy (that depends on the binding energy difference).

Next, we measured efficiencies for agonists that were not studied previously by CRCs (Fig. 4). Choline (Cho) has 2 methylenes between its quaternary nitrogen and hydroxyl (OH) group versus 3 and 4 for 3OH-BTMA and 4OH-PTMA. Cho is a low-affinity, low-efficacy agonist (24) that K_dC_ and K_dO_ values estimated by modeling single-channel kinetics at the human α-ε site indicate has a similar efficiency as does ACh (10). Simulations of binding site structures suggest that an H-bond between the terminal OH and the backbone carbonyl of αW149 serves to position the charged nitrogen of choline away from the aromatic rings that line the cavity, to reduce favorable binding energy (25, 26). Inserting additional methylenes reduces the probability of this H-bond and allows a more-optimal position that increases binding energy relative to Cho.

The CRCs and associated P_O^max^_ and EC_50_ values for 3OH-BTMA and 4OH-PTMA are shown in Fig. 4A left. From the calculated equilibrium dissociation constants, we estimate η is 50% for both agonists (Table 1). Despite the substantial range in affinity and efficacy afforded by the different H-bond propensities, all three of the choline agonists have the same efficiency that is similar to the efficiencies of the agonists shown in Fig. 3. The similarity in the C versus O binding energy *ratio* (but not the difference) for these 3 ligands suggests that the effect of the H-bond on the position of the nitrogen atom applies equally to C and O binding cavities.

Fig. 4A left also shows the CRC of dimethylphenylpiperazinium (DMPP), a nicotinic receptor agonist that is selective for the α3β4 (ganglionic) subtype (27). The result was ηDMPP=52% (Table 1).

Overall, the mean+s.d efficiency calculated from CRCs for the 11 agonists described so far (ACh, Nor, CCh, Ana, TMA, DMP, DMT, Cho, 4OH-BTMA, 4OH-PTMA, DMPP) is 52+2 %. For this entire group of ligands, that covers a huge range in potency and efficacy, the binding energy ratio logK_dC_/logK_dO_ is 0.48. Hence, for all of these agonists binding energy increases by a factor of 2.1 (the inverse of this ratio) when the liganded sites switch from low-to high-affinity at the beginning of the global, channel-opening transition.

Fig. 4A middle shows CRCs for 3 agonists that have a bridge in their head-group. The efficiency values for epibatidine, epiboxidine and cytisine calculated from P_O^max^_ and EC_50_ were η_ebt_=41%, η_ebx_ =46% and η_cyt_=40%. The first two values are similar those estimated previously by kinetic modeling at the human α1–δ binding site (10).

Fig. 4A right shows CRCs for 3 agonists that have extraordinarily low efficacies and affinities. To study these, we added background mutations that did nothing more than increase E_0_ and, hence, increase P_O^max^_ and left-shift EC_50_ (Eqs. 2–4). After correcting for the effects of the background mutations, from the CRC parameters we estimate that in wt AChRs η_TEA_=40%, η_TMP_=41% and η_var_=35%.

For the group of 6 ligands shown in Fig. 4A middle and right (Ebt, Ebx, Cyt, Var, TEA, TMP) the average efficiency was 41+4% (Fig. 4B). For all these ligands, the binding energy ratio (logK_dC_/logK_dO_) is ~0.60, indicating that binding energy increases by a factor of ~ 1.7 when the sites switch from C to O.

Fig. 5A shows a histogram of efficiency values for 17 agonists estimated from single-channel CRCs plus 3 agonists estimated from single-channel kinetic modeling (Atx, Aza, nicotine; Fig. 4C) (10, 28). A goodness of fit test indicates that a bimodal (sum of two Gaussians) frequency distribution is a better fit than a single Gaussian (F (3,16) = 3.37, P=0.044). The two populations have efficiencies of 51+2% and 40+1% (mean+s.d), which is comparable to the mean efficiencies discussed above.

We also estimated the volumes of the head-group of the agonists (Fig. S5 and Table 1) and plotted these versus efficiency (Fig. 5B). A 2D cluster analysis again shows 2 populations with efficiencies of η_1_=52% (n=12) and η2=41% (n=8) with corresponding volumes of v_1_=70.4±8.8 and v_2_±102.2±17.8 Å3 (centroid±s.d). There is an inverse relationship between agonist efficiency and head-group volume.

### CRCs from whole-cell currents

Single-channel CRCs may offer an accurate method for estimating K_dC_ and E_2_, but CRCs constructed from whole-cell responses are more common. In order to ascertain the extent to which η estimated from whole-cell CRCs might be influenced by slow perfusion (that allows desensitization to reduce some peak amplitudes) and heterogeneous receptor properties, we measured whole-cell current amplitude as a function of concentration using 4 agonists, 3 from the high-efficiency group (ACh, CCh and TMA) and 1 from the low-efficiency group (Ebt).

In whole-cell CRCs with maximums normalized to the response to 300 μM ACh response (Fig. 6A), EC_50_ values were left-shifted compared to those in single-channel CRCs by an amount that increased with agonist efficacy (Table 2, left). For example, the left-shift was more substantial for ACh (12.2 μM versus 43 μM) than for TMA (0.84 mM versus 1.2 mM). An independent whole-cell CRC measurement, also made using an automated patch clamp and adult-type AChRs, was 22 μM for EC_50_ for ACh (29). We attribute this left-shift to desensitization (see Materials and Methods). Despite this error, for all 4 agonists the efficiency values estimated from whole-cell CRCs were only slightly smaller than those estimated from single-channel measurements.

**Figure 6.**
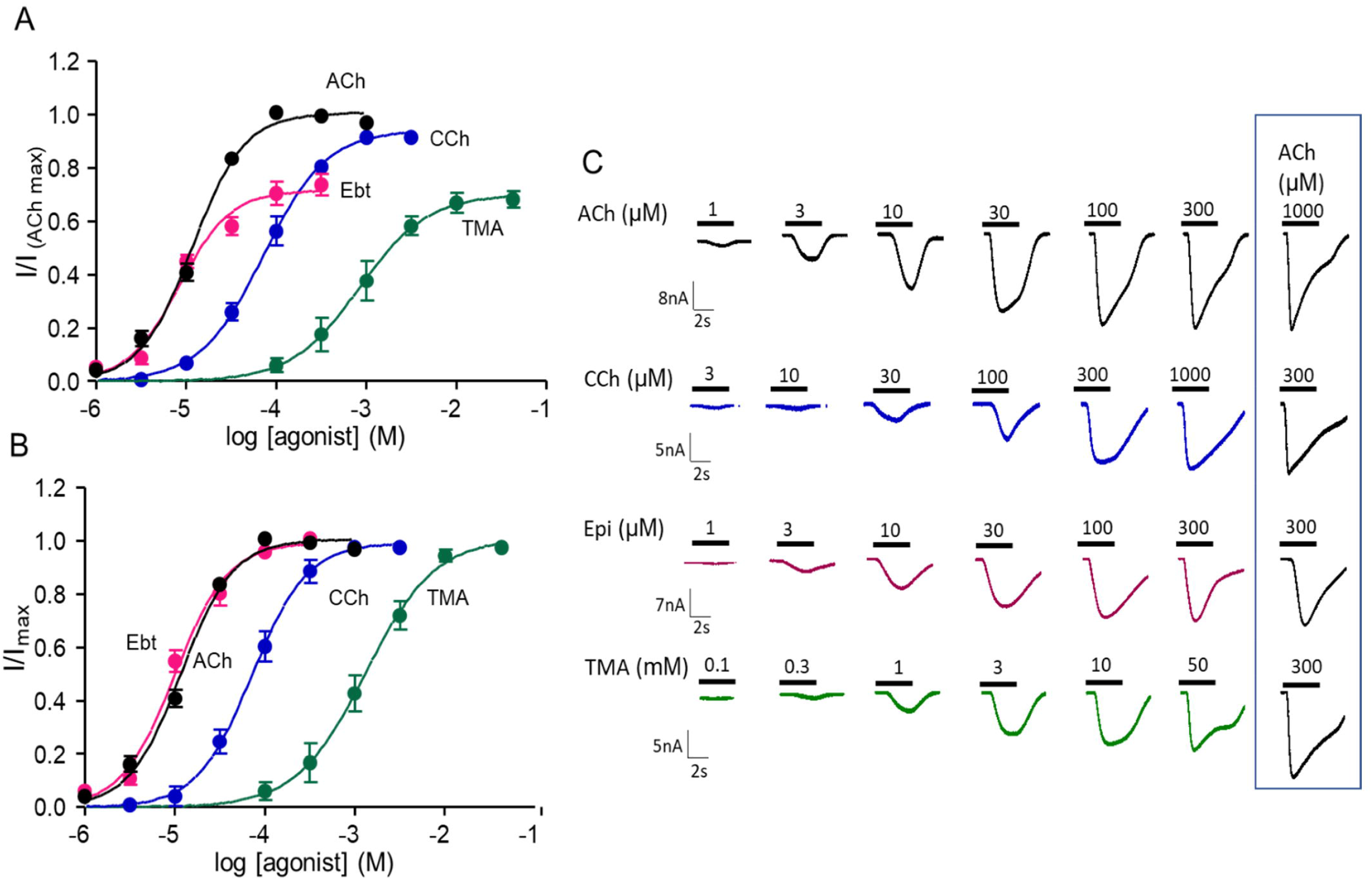
Whole-cell current CRCs. A. Each current response was normalized to that of 300 μM ACh (P_O_=0.93). I^max^ and EC_50_ values in Table 2, left. B. CRCs normalized to I^max^=1. EC_50_ values in Table 2, right. C. Example currents.

**Table 2.** Whole cell CRC parameters and efficiency (η) estimates. Left, EC_50_, I^max^ and n_H_ from CRCs normalized by the response to 300 μM ACh (Fig. 6A; I^max^ for ACh determined from single-channel currents; I^min^, 5.9×10^−7^). Right, EC_50_ from CRCs internally normalized to I^max^=1 (Fig. 6B). n, number of trials (up to 20 cells each).

| Agonist | $I^{\max}$ = response to 300 μM ACh | | | | $I^{\max} = 1$ | n |
| --- | --- | --- | --- | --- | --- | --- |
| | $EC_{50} \pm \text{sem}$<br>(μM) | $I^{\max} \pm \text{sem}$ | $n_H$ | $\eta$ | $EC_{50} \pm \text{sem}$<br>(μM) | |
| ACh | 12.2±0.7 | 0.96 | 1.5 | 0.47 | 12.2±0.7 | 6 |
| CCh | 72.2±11 | 0.91±0.04 | 1.2 | 0.50 | 72.4±6 | 6 |
| TMA | 843±110 | 0.70±0.04 | 1.2 | 0.52 | 1328±160 | 4 |
| Ebt | 8.42±1.0 | 0.72±0.02 | 1.5 | 0.39 | 10±1.2 | 8 |

Because the number of receptors contributing to responses varies from cell to cell and with time, whole-cell CRCs are often normalized so that to the maximum response for each agonist is 1. We did this for the 4 whole-cell CRCs to estimate new values for EC_50_ (Fig. 6B and Table 2, right). It was not possible to estimate efficiency from these plots because information regarding efficacy was removed, but below we show that with knowledge of η and I^min^, I^max^ can be recovered from EC_50_ of a CRC that has been normalized to 1.

### Binding site mutations

K_dC_ and K_dO_ have been measured by kinetic modeling of single-channel currents from mouse, adult-type AChRs having a mutation at one of the 5 aromatic residues at each of the 2 binding sites (30). We calculated from these values η_ACh_ for 21 different mutants (Table S1). Fig. 7A shows that the distribution is Gaussian with η_ACh_=51±4% (mean+s.d.), which is the same as in wt AChRs. The exceptions were αY190A (in loop C) that decreased η_ACh_ to 35%, and mutations of αW149 (in loop B) that increased η_ACh_ by up to 60% in αW149A. Removal of the αY190 side chain results in a ~30% decrease in efficiency, whereas removal of the αW149 side chain results in a ~20% increase in efficiency.

**Figure 7.**
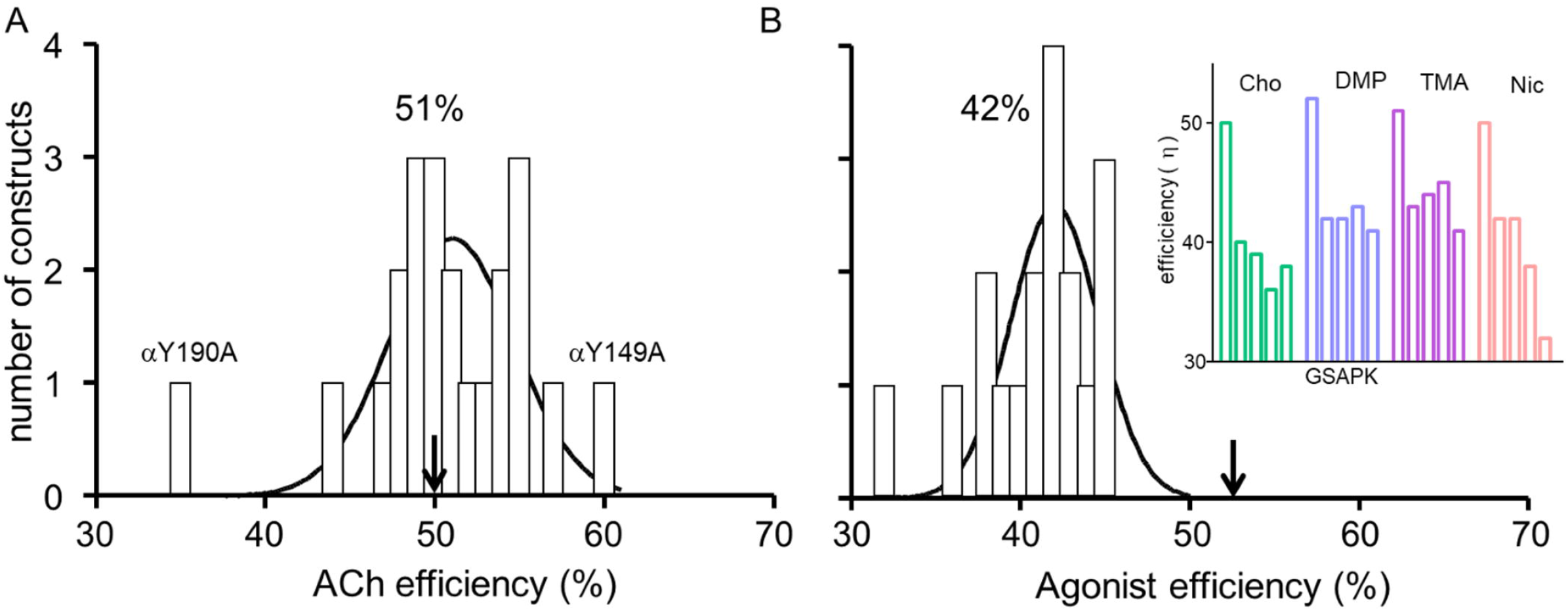
Effect of binding-site mutations on agonist efficiency. Mutations were at both adult-type binding sites. A. Mutations of aromatic amino acids and εP121 (agonist, ACh) (Table S1). Arrow marks wt efficiency. The only substitutions to alter efficiency significantly are αY190A and αW149A. Gaussian fit of frequency distribution for 21 mutants (excluding αY190A) gives η=51±4 % (mean±s.d). B. Mutations of αG153 activated by 4 high-efficiency agonists (Table S2 and inset) (28). Gaussian fit of frequency distribution for all 16 mutation/agonist combinations gives η=42+3 %.

Mutations of a residue on the complementary side of the binding site, εP121, had little effect on ACh efficiency except, perhaps, for the slow-channel myaesthenic syndrome mutation εP121L (31).

Binding and gating equilibrium constants have also been reported for AChRs having a mutation of αG153 (28). This amino acid is in loop B and close to αW149 but does not appear to contact the agonist directly. However, αG153 is interesting because so far it is the only binding site amino acid we know of where mutations decrease K_dC_ (increase binding energy) and increase significantly E_0_. We calculated efficiencies from K_dC_, E_2_ and E_0_ values for 16 different αG153 mutant/agonist combinations using agonists from the high-efficiency population (Table S2).

The distribution of these efficiencies is shown in Fig. 7B. With a αG153 mutation, η 42+3%, which is ~20% smaller than the wt. This is the same efficiency as the low-efficiency agonist population in wt AChRs. αG153 mutations that increase affinity also decrease efficiency. The extent of the reduction in η was similar for all agonists and side chain substitutions, with the exception of αG153K+nicotine. In summary, it appears that a glycine at position α153 allows high efficiency for small-volume agonists that otherwise take on the low efficiency characteristic of large-volume agonists.

### Putting efficiency to use

In this section we show how knowledge of η can simplify and extend CRC analysis. The same efficiency for a group of agonists means that for all, the logK_dC_/logK_dO_ ratio is the same. Hence, the two equilibrium dissociation constants are related by an exponent,

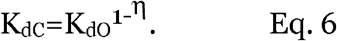

With knowledge of η, only one of the equilibrium dissociation constants needs to be measured. The value of the exponent in Eq. 6 in wt AChRs is ~0.5 for the high-efficiency group of agonists and ~0.6 for the low-efficiency group. Accordingly, Eq. 2 becomes

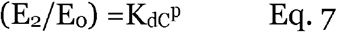

where

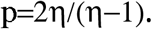

The 2 reflects the number of equivalent binding sites. For the higher efficiency group (η=0.5), p=-2.00 and for the lower efficiency group (η=0.4), p=-1.33.

Taking the log of Eq. 7 and rearranging,

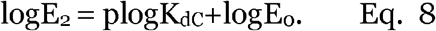

This equation describes an ‘efficiency’ plot, with the x-axis being proportional to agonist binding energy (logK_dC_) and the y-axis being proportional to gating energy (log E_2_). For a group of agonists having the same efficiency, a log-log plot of gating versus binding equilibrium constants is a straight line with a slope (p) that depends on efficiency and a y-intercept that gives the unliganded gating equilibrium constant (Eq. 8). Previously, these constants determined from kinetic modeling were used to estimate average η values for 4 ACh- and Ebt-class agonists at individual AChR binding sites (10). In addition, values of these constants obtained from the literature were fitted by Eq. 8 to estimate E_0_ and average η values for agonists of other receptors, with off-line points reflecting agonists having other efficiencies.

The clustering of AChR efficiency values into 2 populations that correlate with agonist size (Fig. 5) suggests that it may someday be possible to predict approximately an agonist’s efficiency *a priori* from its structure and that of the binding cavity. For example, it is reasonable to guess that in adult-type muscle AChRs other choline or nicotine derivatives will have η~50%, and that congeners of Ebt and TEA will have η~40%. More experiments are needed to test the hypothesis that head-group volume and binding site structure in combination can be used to estimate η. We again note that E_0_ (I^min^) is agonist-independent and needs to be measured only once, so perhaps in the future this important constant will be known for many different receptors.

Given prior knowledge of agonist η and receptor I^min^, the CRC parameters EC_50_ and I^max^ (the response at a single, high agonist concentration) can be estimated from each other, as follows.

First, we calculate EC_50_ from I^max^ (whole-cell CRCs normalized to an ACh response; Table 2, left). The procedure is to solve E_2_ from I^max^ (Eq. 4), then K_dC_ (Eq. 7; η equal to the value shown in Table 2) and then EC_50_ (Eq. 3). Fig. 8B (left) shows that calculated and experimental EC_50_ values are correlated (Pearson’s correlation, r^2^=0.78, P<0.0001). Fig. 8C (left) shows that there is a good match between experimental current amplitudes normalized to an ACh response (Fig. 6A) and those calculated from η according to Eq. 1 using the new EC_50_ estimates. E_2_ has been measured for many AChR agonists (26). Using the above procedure, we estimated corresponding EC_50_ values assuming η=51% and E_0_=7.4×10^−7^ (Table S3). CRCs for these agonists have not been measured, but doing so would test further the ability to use η and E_0_ to calculate EC_50_ from P_O^max^_.

**Figure 8.**
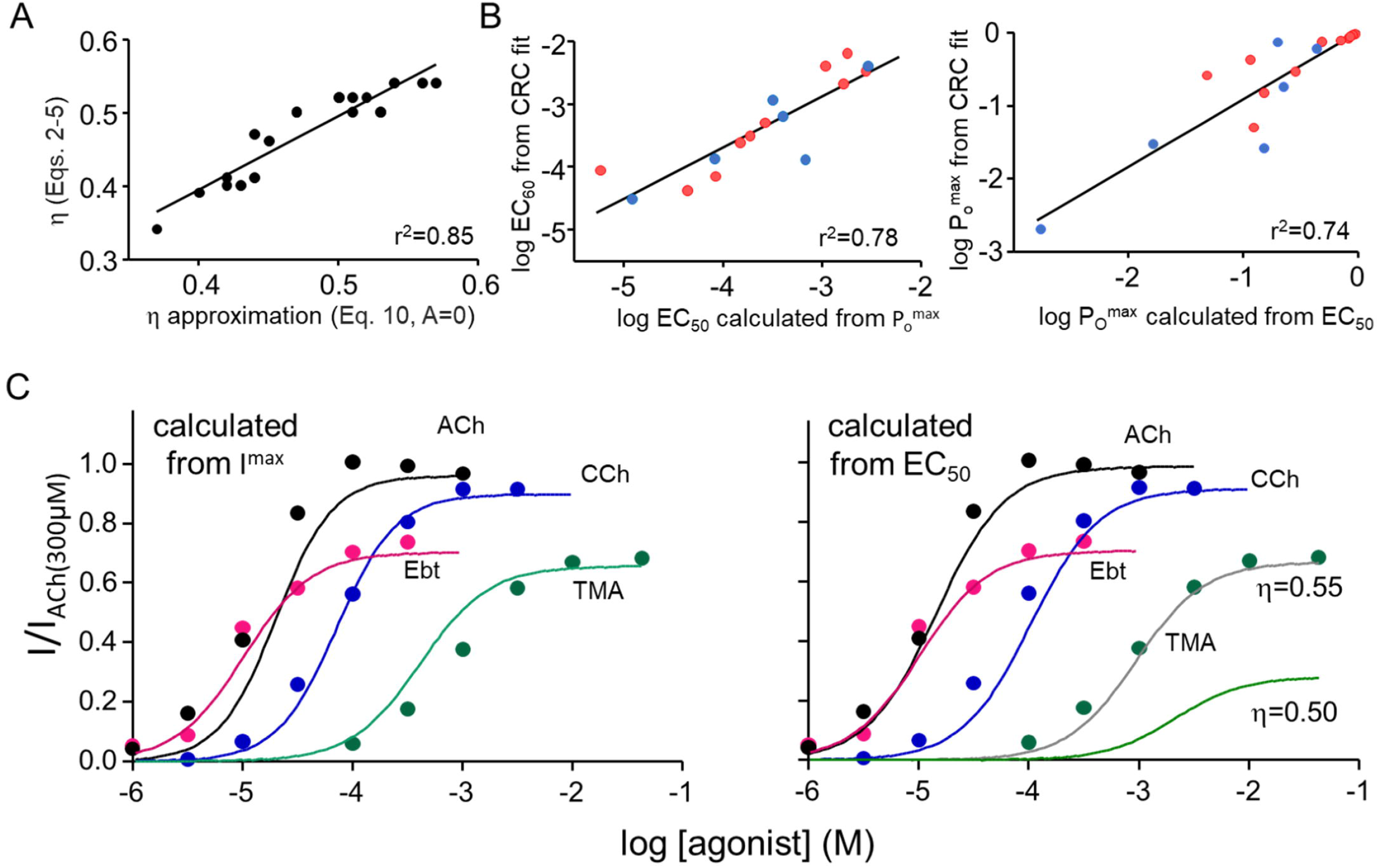
Using efficiency to calculate CRC parameters. A. Approximate efficiencies calculated in a single step (Eq. 10, with A=0) match efficiencies calculated in multiple steps (Eqs.2–5). B. If □ and E_0_ are known, EC_50_ and I^max^ (P_O^max^_) can be calculated from each other. Left, calculated versus measured EC_50_ from single-channel CRCs. η for each agonist given in Table 2. Right, calculated versus measured P_O^max^_ using η=50% (red) or 40% (blue). C. Left, CRCs drawn using calculated EC_50_ and measured I^max^ (Table 2, left) superimposed on whole-cell current responses (Fig. 6A). Right, CRCs drawn using calculated I^max^ and measured EC_50_ (Table 2, right) using η=50% for ACh, CCh and TMA and 40% for Ebt. With TMA, η=55% improves the match.

Second, we calculate I^max^ from EC_50_ values (whole-cell CRCs normalized to 1; Table 2, right). Solving Eqs. 7 and 3 for K_dC_ and setting them equal yields,

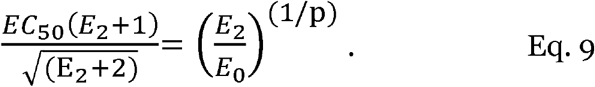

E_2_ and E_0_ can be calculated from I^max^ and I^min^ (Eq. 4). We solved Eq. 9 for I^max^ using known values for η and I^min^ and EC_50_ from normalized CRCs. Fig. 8B (right) is a plot of calculated versus experimental P_O^max^_ values from single-channel CRCs using an approximate value for η, either 50% (ACh, CCh, TMA) or 40% (Ebt). Again, there is rough agreement (Pearson’s correlation, r^2^=0.74, P<0.0001). Next, these calculated I^max^ values were used to generate CRCs (Eq. 1) that were compared to experimental ones that were not normalized to 1 (Fig. 8C, right). The match is good for ACh, CCh and Ebt, but not for TMA. Increasing ηTMA from 0.50 to 0.55 makes the calculated and experimental curves match more closely. The I^max^ value calculated from EC_50_ by using Eq. 9 is sensitive to the value of η (Fig. S6). Nonetheless, knowledge of agonist efficiency allows efficacy information to be recovered approximately from a CRC that has been normalized to a maximum response of 1.

In addition, knowledge of η allows the estimation of E_0_ from a single CRC. E_0_ is of critical importance because it sets the baseline level from which agonists increase P_O_, but it is often small and difficult to measure directly (8, 32, 33). However, a fold-change in E_0_ caused by a mutation or a modulator will produce the same fold-change in E_2_ (Eq. 2) and, hence, a change in both EC_50_ and P_O^max^_ (Eqs. 3 and 4).

The procedure we used to estimate E_0_ (I^min^) from a CRC of a wt receptor is first to solve for E_2_ and K_dC_ from P_O^max^_ and EC_50_ as described above, then solve for E_0_ by using Eq. 8. Fig. S7 shows E_0_ values so-calculated from single-channel CRC parameters. The mean result is reasonably close to the experimentally-determined value (17).

Finally, it is possible to gain an approximate estimate of η from CRC parameters in a single step. Taking the log of Eq. 9 and rearranging,

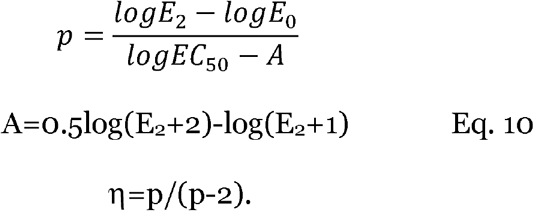

E_2_=I^max^/(1-I^max^) and E_0_=I^min^. For many AChR agonists E_2_<25 and EC_50_<10^−3^ M, so A will usually be much less than log(EC_50_). Hence, a reasonable approximation for agonist efficiency can be obtained simply by using Eq. 10 with A equal to zero. Fig. 8A shows that η-values calculated using this shortcut indeed approximate the more-exact values calculated stepwise using Eq. 2-5.

## Discussion

The notion of agonist efficiency arose from an experimental observation – in neuromuscular AChRs, the binding energy ratio logK_dC_/logK_dO_ is the same for many different nicotinic agonists (23). Later, this ratio was associated with the efficiency at which agonist binding energy is converted into receptor gating energy (10). Here, we show that agonist efficiency can be estimated from the asymptotes and midpoint of a CRC. Below, we discuss the nature and distribution of efficiency values obtained from single-channel and whole-cell CRCs, some structural implications of efficiency, and applications of efficiency to CRC analysis.

### Efficiency

Agonist efficiencies are the same whether obtained from single-channel or wholecell currents, and from a CRC or by detailed kinetic modeling. They are the same in wild-type AChRs that have 2 binding sites or in crippled AChRs that have just 1 operational site. Efficiency values are the same in mouse and human AChRs and with many mutations at the binding sites (exceptions discussed below) or in distant regions that do not affect binding. In AChRs, efficiency is a robust agonist attribute. At adult neuromuscular synapses, half of the available neurotransmitter binding energy is converted into kinetic energy of the channel-opening conformational change.

In AChRs there are 2 populations of η-values, at 51±2% and 40±4% (Fig. 5A). Despite the small standard deviations, we suspect that the variance within each group arises from actual, ligand-specific differences rather than from measurement errors, for the following reasons, i) Efficiency is a ratio of logarithms and therefore is not sensitive to errors in the measured values of EC_50_ and P_O^max^_. For example, changing EC_50_ or P_O^max^_ (Table 1) by +10% changes the calculated η value by <1%. ii) The order of η-values within the high efficiency group is the same in single-channel and whole-cell experiments (TMA>CCh>ACh). iii) A small difference in efficiency leads to a large difference in efficacy calculated from EC_50_ (Fig. S6). The single-channel η-value for TMA predicts the experimental, whole-cell CRC more-accurately than does the group value (Fig. 8C, right). We hypothesize that the efficiency difference between, for example, ACh and TMA is meaningful (Fig. 3B).

The observation that mutations of αG153 shift η for 4 agonists from the high-to the low-efficiency population supports the existence of two discrete η populations. Although the distribution of agonist efficiency appears to be modal rather than continuous, the high accuracy of experimental η estimates must be considered. More experiments might reveal if other η populations exist or if small differences between agonists or mutations are meaningful. For example, the observations that η is modestly lower with εP121 substitutions (Table S1), higher with most non-aromatic substitutions of αY198 (Table S1) and usually lowest with a K substitution at αG143 (Table S2) might prove to be meaningful. Likewise, experiments might show that the 35% efficiency values for varenicline and ACh+αY190A indicate the existence of a third population.

### Structural implications

That a group of agonists have the same efficiency means that all members have the same binding energy ratio, logK_dC_/logK_dO_ (Eq. 5). Below, we discuss implications of this result with regards to i) rearrangements at the binding site, ii) agonist volume, iii) the bimodal efficiency distribution and iv) binding-site mutations.

The energy of low-affinity binding is proportional to logK_dC_ and is determined mainly not by diffusion but rather by a local rearrangement at the binding site called ‘catch’. The energy of the switch from low-to high-affinity is proportional to (logK_dO_-logK_dC_), occurs at the beginning of the global, channel-opening isomerization and is called ‘hold’ (30, 34). As discussed elsewhere (34), ‘hold’ is related to, and possibly the same as, an intermediate (pre-opening) gating state called ‘flip’ that has been detected directly (35, 36). ‘Flip’ refers to a brief shut state that is high affinity, and ‘hold’ refers to the rearrangement of the binding site that generates such a state (37). Regardless, a group efficiency implies that for all members, the energy change in hold is 1/(1-η) times that of catch. This factor is ~2 for high- and ~1.7 for low-efficiency agonists.

This linear relationship between catch and hold energy changes suggests that the associated structural changes, too, are related. Accordingly, the observation that many agonists have the same efficiency suggests that the binding-site rearrangements in catch and in hold can be considered as two stages of a single conformational sweep. Although ‘binding’ and ‘gating’ have long been considered to be distinct processes (38), a group efficiency implies that they are components of a multipart structural-change cascade. In AChRs, this cascade begins with a ‘touch’ by the agonist that takes place after the ligand has diffused to its target but before binding site rearrangements that form the low-affinity complex, and ends when ions begin to cross the membrane. The ‘catch-and-hold’ sweep of the binding sites is the first part of this cascade, with η quantifying the strength of the connection between the binding (catch) and gating (hold) components.

It remains to be determined whether or not a shared binding energy ratio for a group of related agonists is a general feature of receptor activation. It appears that in some receptors other than neuromuscular AChRs there is a linear relationship between log gating and log binding equilibrium constants for related ligands, and that association to C is slower than diffusion. These results raise the possibility that a shared logK_dC_/logK_dO_ ratio and a correlation between structural changes in low- and high-affinity complex formation are not exclusive to AChRs (10).

Members of both efficiency populations can have a quaternary amine (TMA, TEA) or a secondary amine (DMPP, Ebt). Hence, it does not appear that the bimodal distribution in efficiency reflects this aspect of the agonist’s head group. The inverse correlation between agonist head-group volume and efficiency is more relevant (Fig. 5B). Simulations of AChR structures suggest that the agonist binding cavity is smaller in O compared to in C and, hence, that binding site contraction is a structural correlate of ‘hold’ (25). In addition, kinetic analyses of AChR gating indicate that in the channel-opening isomerization, ‘hold’ is followed by a rearrangement of the extracellular domain (4, 34, 39, 40).

These results lead to hypothesize that large-volume, low-efficiency agonists encounter steric hindrance when the binding cavity contracts in ‘hold’, to limit the shrinkage and, hence, the mechanical force applied to the next element in the gating sequence, the extracellular domain. We imagine that small, high-efficiency ligands fit comfortably into both C and O pockets but that large, low-efficiency agonists do not fit easily into the smaller O cavity and so support a smaller contraction. According to this hypothesis, large ligands transfer less energy to the next step in the conformational cascade and thus have low efficiencies.

In support of this idea, the smallest agonists we tested, DMP, DMT and TMA, have the largest efficiencies (Fig. 3B). Further, simulations show that compared to ACh, the binding cavity is smaller with TMA and the extent of cavity contraction is smaller with the low-efficiency agonist Ebx (25). However, the relationship between agonist volume and efficiency is not simple because Ebt and TEA have similar efficiencies despite a substantial difference in volume (Table 1).

There are 2 efficiency populations (Fig. 5A). One possible explanation is that each efficiency group reflects a different ‘hold’ binding site conformation. In this view, the high-affinity cavity can adopt only a limited number (so far, 2) of ‘preset’ structures and is not malleable or able to adapt its shape to each agonist. Perhaps small versus large agonists allow the pocket to adopt alternative contracted shapes, with all agonists larger than some threshold forcing the less-efficient shape. Another hypothesis for the bimodal distribution of η is that there are two discrete energy transfer pathways that supply energy for activating the extracellular domain. Both paths are activated with smaller agonists but one (or both) is compromised when the pocket is ‘stretched’ by a large ligand. Both of these hypotheses are speculations that can be tested experimentally.

Aromatic side chains at the binding site govern agonist affinity. While most mutations of these have little or no effect on η, another clue regarding the structural basis of efficiency is that the efficiency of ACh is reduced by 30% by the mutation αY190A (in loop C) and increased by 20% by the mutation αW149A (in loop B). αY190 appears to be the most-important aromatic side chain with regards to the propagation of structural changes from the binding site in channel opening (22, 41). That the mutations αY190F and αY190W have little effect on ACh efficiency suggests that the key interaction here is with the aromatic ring rather than with the OH group.

All 4 mutations of αG153 (in loop B) reduced the efficiency of all 4 tested agonists. Again, the drop appeared to be modal, reducing the average efficiency from 51% to 42%. At this juncture we do not have a hypothesis for the structural basis for this decrease in efficiency. Perhaps molecular dynamics simulations can test if flexibility of the loop B backbone promotes high efficiency. It will be worthwhile to ascertain experimentally the extent to which the αW149A and αG153 mutations are correlated.

Efficiency estimates for the two populations are the same whether measured in whole receptors (that have α-δ and α-ε binding sites) or in receptors having only one functional site (10). This indicates that the energy changes that contribute to efficiency are determined mainly by local ligand-protein interactions at each binding site, with little or no energy transfer between sites. The efficiency of Ebx is somewhat higher in whole receptors compared to at α-δ alone so it is possible that the efficiency of this agonist is modestly greater at one site (α-ε) compared to the other. Agonist affinity is greatest at the fetal (α1–γ) neurotransmitter binding sites (56% for ACh) and it is possible that the some of the small, agonist-dependent differences in η between adult sites are meaningful.

### Applications

Because η values are modal, efficiency can be used to classify agonists. Someday, efficiency may stand alongside affinity and efficacy as a core agonist attribute. An approximate value for agonist efficiency can be calculated from CRC parameters by using Eq. 10, with A =0 (Fig. 8A). To make the calculation even easier, Eq. 10 can be rearranged to express η directly in terms of CRC parameters,

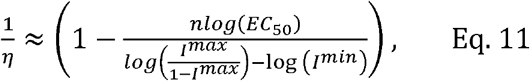

where n is the number of agonist binding sites and is proportional to n_H_ (42). Eq. 11 offers an easy way to estimate approximately an agonist’s efficiency from a CRC. If I^min^ is not known, it may be possible to compare efficiencies of different agonists by using a common value, for instance 10^−6^.

Knowing η is useful because it allows EC_50_ and I^max^ to be estimated from each other. With knowledge of η, EC_50_ and an entire CRC can be calculated knowing only the responses at the low- and high-concentration asymptotes (Fig. 8C). Given η and I^min^, the response at just one agonist concentration, that which produces I^max^, needs to be measured in order to estimate EC_50_. Having this ability could facilitate drug screening.

It is common practice in CRCs to normalize I^max^ to 1 and lose information regarding agonist efficacy. We have shown that given prior knowledge of η and I^min^, and an experimental estimate of EC_50_, Eq. 9 can be solved numerically for E_2_ and, hence, I^max^. The ability to compute an absolute CRC from a normalized one could be useful once the values of the agonist’s efficiency and the receptor’s constitutive activity are established. The main caveat is that the calculated efficacy is very sensitive to the value of η.

## Author contributions

Dinesh C. Indurthi: designed and performed research; analyzed data.

Anthony Auerbach: conceptualized and designed research; analyzed data and wrote the manuscript.

## Acknowledgements

We thank Mary Teeling, Marlene Shero and Janet Jordan for technical assistance and Pablo M. Paez for allowing access to his IonFlux. Supported by NS-064969 and GM-121463

## Competing interests

The authors declare no competing financial or non-financial interests.

## Notes

### Competing Interest Statement

The authors have declared no competing interest.

### Summary of Updates

added: 1. efficiency calculated from whole-cell currents 2. efficiency of alphaG153 mutations 3. using efficiency to calculate efficacy (Imax) from potency (EC50).

