## Supplemental Information for "Agonist efficiency from concentration-response curves: structural implications and applications"

running title: agonist efficiency

Dinesh C. Indurthi and Anthony Auerbach‡

Department of Physiology and Biophysics

State University of New York at Buffalo

Buffalo NY 14214

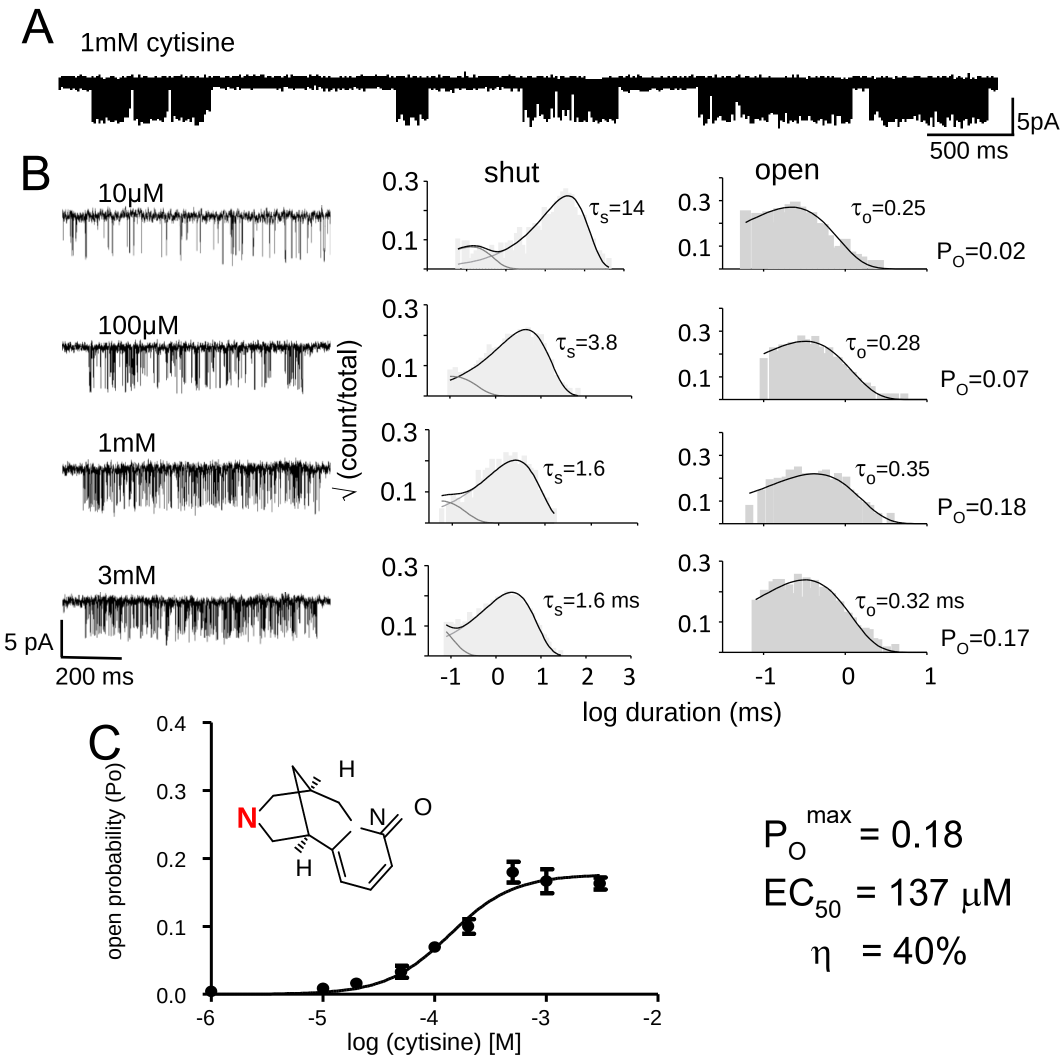


**Figure S1.** Constructing a CRC from single-channel currents (agonist, cytisine). A. Low-resolution view of cell-attached, single-channel currents showing clusters of openings and closing arising from a single AChR separated by long silent periods in which all AChRs in the patch are desensitized (open down). B. High-resolution views of example clusters at different cytisine concentrations and corresponding intra-cluster interval-duration histograms. The predominant shut and open interval durations ($\tau_S$ and $\tau_O$) were used to calculate an open-channel probability ($P_O$) at each agonist concentration. C. CRC fitted by Eq. 1 to estimate $EC_{50}$ and $P_O^{max}$ (symbols, mean±s.e.m.). Efficiency ($\eta$) was calculated by using Eqs. 2-5.

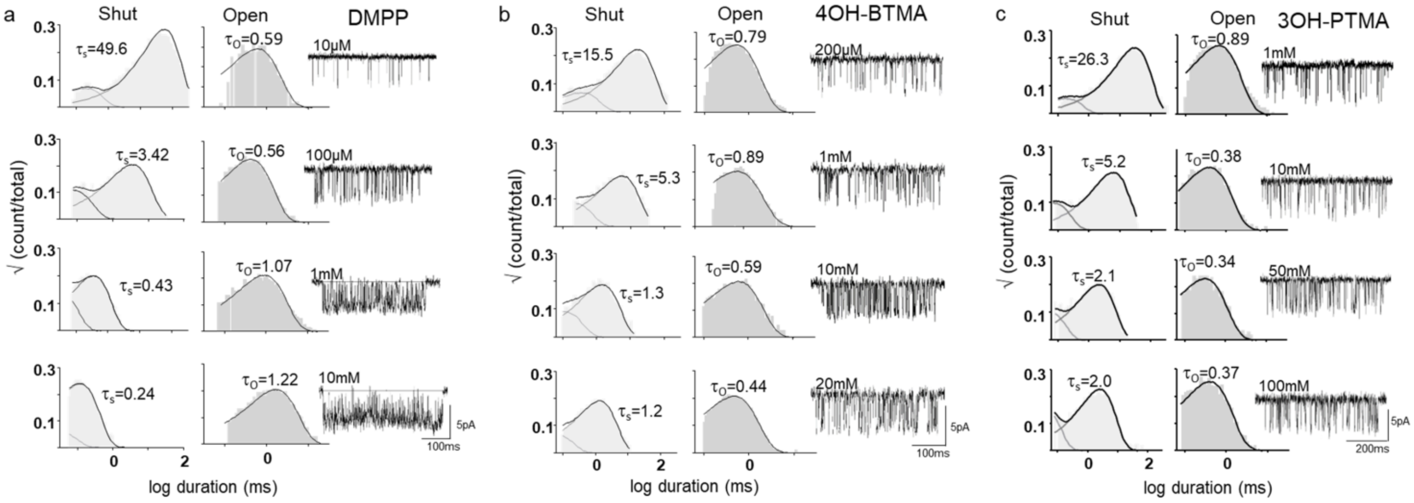


**Figure S2.** Activation of adult-type AChRs by agonists (open in down). Agonist abbreviations in Materials and Methods. Interval-duration histograms and an example cluster. The predominant intra-cluster shut and open interval durations ($\tau_s$ and $\tau_o$) are in ms. CRCs are shown in Fig. 4A.

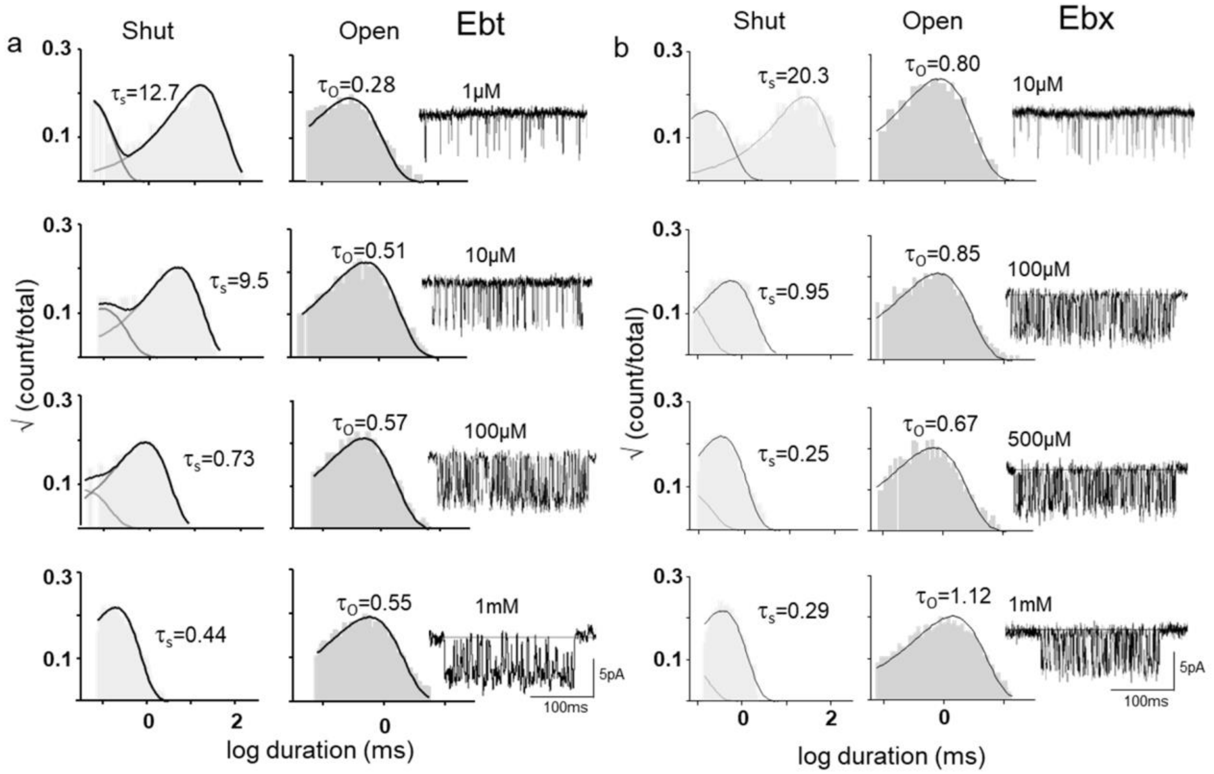


**Figure S3.** Activation of adult-type AChR in the presence of Ebt (epibatidine) and Ebx (epiboxidine). Time constants are ms. CRCs are shown in Fig. 4B.

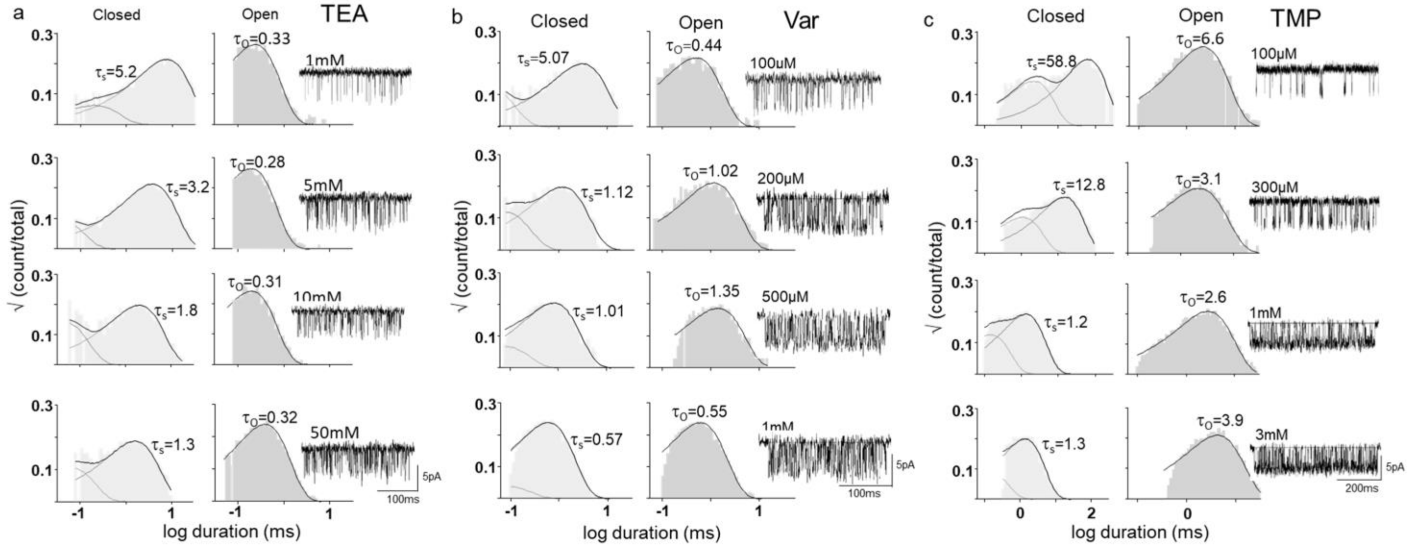


**Figure S4.** Activation of adult-type AChR by TEA (tetraethylammonium), Var (varenicline) and TMP (tetramethylphosphonium). Time constants are ms. CRCs are shown in Fig. 4C.

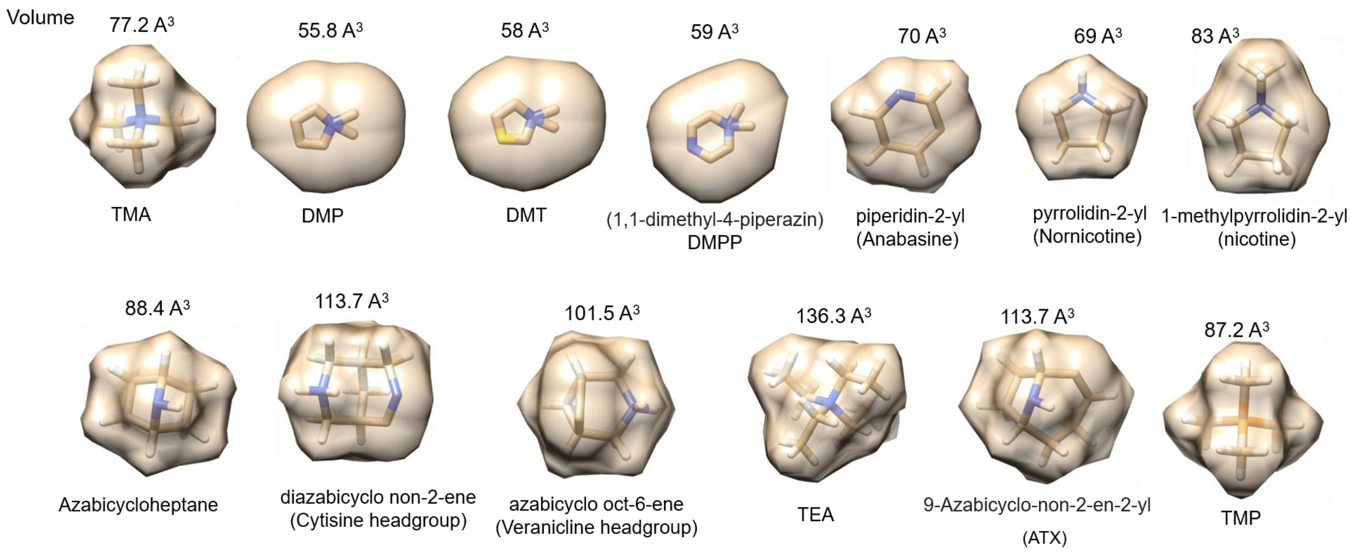


**Figure S5.** Agonist head-group volume. Not shown: the head-groups of ACh, CCh, choline, 3OH-P and 4OH-B are the same as TMA, and those of Ebt and Ebx are the same as azabicycloheptane.

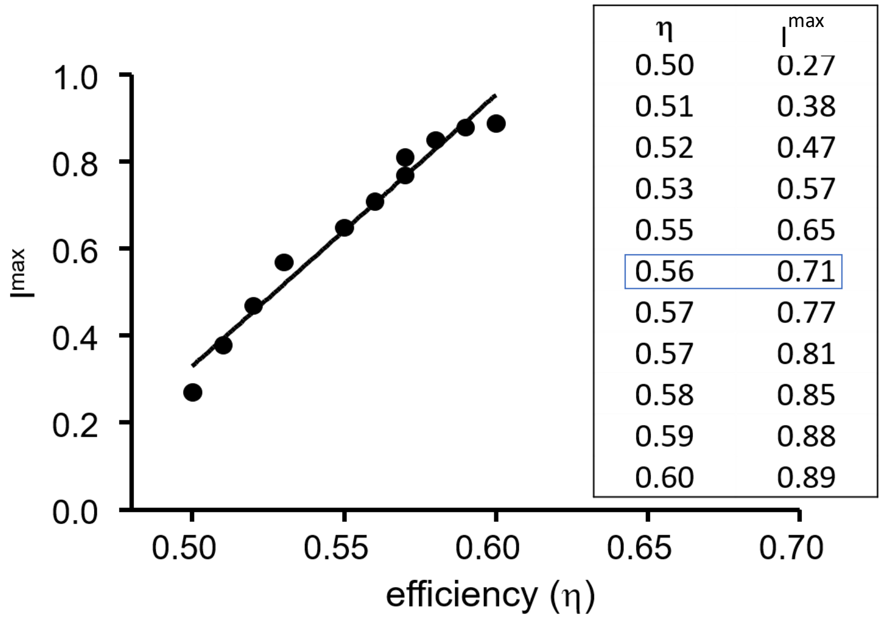


**Figure S6.** Using efficiency (η) to calculate $I^{max}$ from $EC_{50}$ (from whole cell CRCs normalized to a maximum of 1; Table 2, right) (see Fig. 8B). Agonist, TMA; experimental $EC_{50}$, 1.3 mM; $E_0$=5.2x10$^{-7}$. From non-normalized CRCs, the true $I^{max}$= 0.70 (Table 2, left). $I^{max}$ calculated by using Eq. 9 is very sensitive to η, making it possible to estimate an agonist's efficiency by matching its calculated and experimental (non-normalized) efficacies.

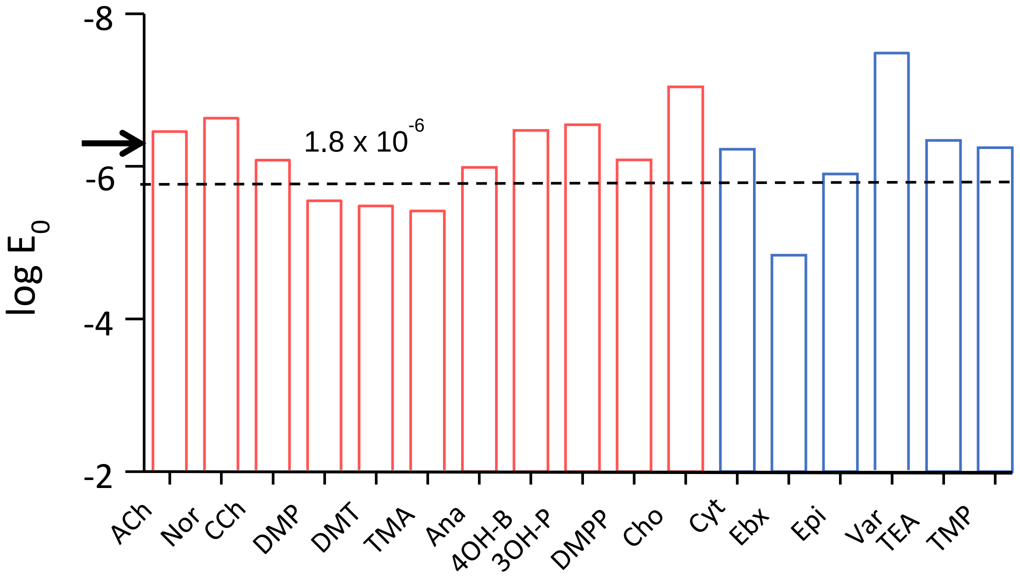


**Figure S7.** Unliganded gating equilibrium constant ($E_0$) estimated from CRC parameters. The assumed η values were either 52% (red) or 41% (blue). The calculated mean $E_0$ of 1.8 x $10^{-6}$ (dashed line) is approximately the same regardless of η and reasonably close to the $E_0$ value estimated by using other methods, 7.4 x $10^{-7}$ (arrow).

**Table S1. ACh efficiency after mutation of a binding site residue (aromatics and εP121).**

| | $K_{dC}$ (mM) | $K_{dO}$ (μM) | log $K_{dC}$ | log $K_{dO}$ | η |
|---|---|---|---|---|---|
| WT | 0.166 | 0.028 | -3.78 | -7.55 | *0.50* |
| αY93W | 2.01 | 1.20 | -2.70 | -5.92 | *0.54* |
| H | 3.20 | 2.90 | -2.49 | -5.54 | *0.55* |
| A | 1.21 | 1.30 | -2.92 | -5.89 | *0.50* |
| F | 2.59 | 5.80 | -2.59 | -5.24 | *0.51* |
| S | 6.24 | 14.0 | -2.20 | -4.85 | *0.55* |
| αW149Y | 2.41 | 3.00 | -2.62 | -5.52 | *0.53* |
| F | 12.8 | 19.0 | -1.89 | -4.72 | *0.60* |
| A | 28.8 | 260.0 | -1.54 | -3.59 | *0.57* |
| αY190F | 3.60 | 16.0 | -2.44 | -4.80 | *0.49* |
| W | 6.46 | 56.0 | -2.19 | -4.25 | *0.48* |
| A | 16.5 | 1900 | -1.78 | -2.72 | *0.35* |
| αY198F | 0.23 | 0.053 | -3.64 | -7.28 | *0.50* |
| H | 5.70 | 9.10 | -2.24 | -5.04 | *0.55* |
| W | 0.61 | 0.88 | -3.21 | -6.06 | *0.47* |
| S | 3.90 | 12.0 | -2.41 | -4.92 | *0.51* |
| T | 9.20 | 38.0 | -2.04 | -4.42 | *0.54* |
| L | 4.10 | 21.0 | -2.39 | -4.68 | *0.49* |
| A | 7.50 | 41.0 | -2.12 | -4.39 | *0.52* |
| εP121L | 0.72 | 2.20 | -3.14 | -5.66 | *0.44* |
| Y | 1.27 | 3.00 | -2.90 | -5.52 | *0.48* |
| G | 1.00 | 1.20 | -3.00 | -5.92 | *0.49* |

Equilibrium dissociation constants ($K_d$) of ACh to resting (C) and active (O) conformations of the binding site (Fig. 1) estimated by kinetic modeling (1). Efficiency (η) is 1-log$K_{dC}$/log$K_{dO}$.

**Table S2. Efficiency after mutation os αG153.**

| | Ligand | $E_2$ | $E_0$ | $K_{dc}$ | $K_{dO}$ | log $K_{dc}$ | log $K_{dO}$ | η |
|---|---|---|---|---|---|---|---|---|
| αG153 | Cho | 0.05 | 7.4E-07 | 4.0E-03 | 1.5E-05 | -2.40 | -4.81 | ***0.50*** |
| S | Cho | 0.63 | 1.9E-05 | 3.7E-04 | 2.0E-06 | -3.43 | -5.69 | ***0.40*** |
| A | Cho | 1.18 | 4.2E-05 | 2.9E-04 | 1.7E-06 | -3.54 | -5.77 | ***0.39*** |
| P | Cho | 1.1 | 4.8E-05 | 1.5E-04 | 1.0E-06 | -3.81 | -5.99 | ***0.36*** |
| K | Cho | 3 | 1.4E-04 | 2.6E-04 | 1.8E-06 | -3.59 | -5.75 | ***0.38*** |
| αG153 | DMP | 0.4 | 7.4E-07 | 2.1E-03 | 2.9E-06 | -2.68 | -5.54 | ***0.52*** |
| S | DMP | 6.14 | 1.9E-05 | 1.8E-04 | 3.2E-07 | -3.74 | -6.50 | ***0.42*** |
| A | DMP | 12.2 | 4.2E-05 | 1.9E-04 | 3.6E-07 | -3.71 | -6.44 | ***0.42*** |
| P | DMP | 13.8 | 4.8E-05 | 2.0E-04 | 3.8E-07 | -3.69 | -6.42 | ***0.43*** |
| K | DMP | 20.6 | 1.4E-04 | 2.3E-04 | 5.9E-07 | -3.64 | -6.23 | ***0.41*** |
| αG153 | TMA | 2.5 | 7.4E-07 | 8.1E-04 | 4.4E-07 | -3.09 | -6.36 | ***0.51*** |
| S | TMA | 25.8 | 1.9E-05 | 8.1E-05 | 7.0E-08 | -4.09 | -7.16 | ***0.43*** |
| A | TMA | 285 | 4.2E-05 | 3.9E-05 | 1.5E-08 | -4.41 | -7.82 | ***0.44*** |
| P | TMA | 179 | 4.8E-05 | 1.0E-04 | 5.2E-08 | -4.00 | -7.29 | ***0.45*** |
| K | TMA | 627 | 1.4E-04 | 1.7E-05 | 8.1E-09 | -4.76 | -8.09 | ***0.41*** |
| αG153 | Nicotine | 0.87 | 7.4E-07 | 1.0E-03 | 9.2E-07 | -3.00 | -6.04 | ***0.50*** |
| S | Nicotine | 12.34 | 1.9E-05 | 9.2E-05 | 1.1E-07 | -4.04 | -6.94 | ***0.42*** |
| A | Nicotine | 17.7 | 4.2E-05 | 1.1E-04 | 1.8E-07 | -3.94 | -6.76 | ***0.42*** |
| P | Nicotine | 12.5 | 4.8E-05 | 3.8E-05 | 7.4E-08 | -4.42 | -7.13 | ***0.38*** |
| K | Nicotine | 549 | 1.4E-04 | 1.2E-07 | 6.1E-11 | -6.92 | -10.22 | ***0.32*** |
| E | Nicotine | 32.05 | 4.5E-05 | 2.6E-05 | 4.0E-09 | -4.59 | -8.40 | ***0.45*** |
| R | Nicotine | 23.64 | 3.8E-05 | 3.2E-05 | 5.7E-09 | -4.49 | -8.25 | ***0.45*** |

Equilibrium dissociation constants ($K_d$) of 4 agonists to resting (C) and active (O) conformations (Fig. 1) estimated by kinetic modeling (2). Efficiency (η) is 1-$\log K_{dC}/\log K_{dO}$.

**Table S3. $EC_{50}$ from efficacy given η and $E_0$ for analogues of choline.**

| Input values | | | | Calculated values | | | | | |
|---|---|---|---|---|---|---|---|---|---|
| **Ligand** | **$E_2$** | **η** | **$E_0$** | **$P_O^{max}$** | **log $E_0$** | **log $E_2$** | **log $E_2$-log$E_0$** | **$K_{dC}$ (mM)** | **$EC_{50}$ (mM)** |
| **TMA** | 2.54 | 0.52 | 5.2E-7 | 0.72 | -6.28 | 0.40 | 6.69 | 0.82 | 0.49 |
| **ETMA** | 0.25 | 0.52 | 5.2E-7 | 0.20 | -6.28 | -0.60 | 5.68 | 2.39 | 2.86 |
| **PTMA** | 0.29 | 0.52 | 5.2E-7 | 0.22 | -6.28 | -0.54 | 5.75 | 2.61 | 2.61 |
| **BTMA** | 2.44 | 0.52 | 5.2E-7 | 0.71 | -6.28 | 0.39 | 6.67 | 0.83 | 0.51 |
| **Cho** | 0.05 | 0.52 | 5.2E-7 | 0.05 | -6.28 | -1.30 | 4.98 | 5.01 | 6.84 |
| **3OH-PT** | 0.15 | 0.52 | 5.2E-7 | 0.13 | -6.28 | -0.82 | 5.46 | 3.02 | 3.85 |
| **4OH-B** | 0.71 | 0.52 | 5.2E-7 | 0.42 | -6.28 | -0.15 | 6.14 | 1.47 | 1.42 |
| **Cl Cho** | 0.19 | 0.52 | 5.2E-7 | 0.16 | -6.28 | -0.72 | 5.56 | 2.71 | 3.37 |
| **2OH-P** | 0.02 | 0.52 | 5.2E-7 | 0.02 | -6.28 | -1.70 | 4.59 | 7.65 | 10.7 |
| **Cholamine (pH 9.0)** | 0.04 | 0.52 | 5.2E-7 | 0.04 | -6.28 | -1.40 | 4.89 | 5.56 | 7.63 |

Using η and $E_0$ to calculate $EC_{50}$ and $K_{dC}$ (Eq. 7) from $E_2$ (3). TMA (tetramethyl ammonium), ETMA (ethyltrimethyl ammonium), PTMA (propyltrimethyl ammonium), BTMA (butyltrimethyl ammonium), Cho (choline), 3OH-PTMA (3-hydroxypropyl trimethyl ammonium), 4OH-BTMA (4-hydroxybutyl trimethyl ammonium), Cl Cho (2-chloroethyl trimethyl ammonium), 2OH-PTMA (2-hydroxypropyl trimethyl ammonium), cholamine (2-aminoethyl trimethyl ammonium).
